# Distinct Ca^2+^ signatures of leptomeningeal fibroblast subgroups in awake mouse brains in health and inflammation

**DOI:** 10.1101/2025.05.13.653681

**Authors:** Chen He, Søren Grubb, Lechan Tao, Kıvılcım Kılıç, Xiao Zhang, Anna Devor, Neha Zahoor Sheikh, Changsi Cai

## Abstract

The leptomeninges, composed of the arachnoid mater, and pia mater, contains distinct subgroups of fibroblasts that differ in location and transcriptomic profiles. These fibroblasts contribute to the blood–cerebrospinal fluid barrier under physiological conditions, participate in fibrosis, and support blood–brain barrier integrity during injury and disease. However, their Ca^2+^ signaling profiles and underlying mechanisms in health and disease remain poorly understood. In this study, we divided leptomeningeal fibroblasts into three subgroups based on their locations: arachnoid fibroblasts, pia mater fibroblasts and perivascular fibroblasts. We employed two-photon microscopy in awake transgenic mice expressing Ca²⁺ indicators in leptomeningeal fibroblasts to investigate spontaneous and behaviorally evoked Ca²⁺ transients across different fibroblast subgroups. We found that each subgroup exhibits a distinct Ca²⁺ activity profile, with pia mater fibroblasts showing the highest-amplitude Ca²⁺ transients. Moreover, these fibroblasts displayed unique responses to both whisker air-puff stimulation and locomotion. We further demonstrated, using a chronically implanted cannula beneath the cranial window, that locomotion-associated vasodilation is followed by TRPV4 channel-mediated fibroblast Ca²⁺ elevations. Finally, systemic inflammation induced by lipopolysaccharide (LPS) reduced spontaneous Ca²⁺ transients in pia mater fibroblasts, likely due to macrophage infiltration following the inflammatory response. For the first time, this study characterizes spontaneous and behaviorally evoked Ca²⁺ dynamics in distinct leptomeningeal fibroblast subgroups in awake animals, providing novel insights into the functional roles of leptomeningeal fibroblasts in the healthy and diseased brain.

## Introduction

The meninges are protective membranes surrounding the brain, providing physical support against external mechanical shocks, facilitating cerebrospinal fluid circulation, and offering immunological protection by responding to infections or injury [1, 2]. The meninges comprise three layers - the dura mater, arachnoid mater, and pia mater - and consist of, e.g., extracellular matrix, fibroblasts (FBs), immune cells, endothelial cells of blood vessels, and vascular mural cells. The arachnoid mater and the pia mater, are collectively known as the leptomeninges. The term “leptomeninges” comes from the Greek word “leptós,” which means thin, reflecting the delicate nature of these membranes. The locations and functions of FBs have recently attracted increasing attention but are still in the initial phase of understanding [2-5]. Advances in methodologies, including sophisticated lineage tracing techniques and single-cell RNA sequencing, have significantly enhanced our understanding of FBs, which reside in all sublayers of leptomeninges [2, 6]. FBs in the arachnoid mater, pia mater, and perivascular space display region-specific molecular profiles [2, 7], developmental trajectories [8, 9], suggesting they have distinct yet incompletely understood functional roles in maintaining the central nervous system (CNS) in health and disease.

FBs within the arachnoid mater (ArachF) form the arachnoid barrier, a critical component regulating cerebrospinal fluid homeostasis and maintaining CNS immune privilege [10]. They also form a reticular cell layer that expresses the lymphatic marker, *Prox-1*, and forms a collagen matrix which adheres the arachnoid mater to the pial surface via collagen fibers crossing the subarachnoid space [3-5]. FBs lining the subarachnoid space share similar gene markers [2]. They are strategically positioned to interact closely with cerebrospinal fluid and produce extracellular matrix (ECM) proteins that structurally support and organize the fluid-containing spaces, and contribute to the structural integrity of the leptomeningeal-cerebral boundary [11]. Perivascular fibroblasts (PVF) directly adhere to cerebral blood vessels. PVFs surround large-diameter arterioles and venules, positioning them uniquely to support vascular structures, potentially influencing neurovascular development and regulating vascular permeability [8, 12, 13]. However, the precise physiological functions of leptomeningeal FBs remain poorly understood.

Previous research has explored the roles of brain FBs in CNS pathologies. Following acute injury, such as stroke or traumatic brain injury, local FBs rapidly proliferate and contribute extensively to fibrosis and scar formation [14]. Additionally, there is growing evidence indicating that FBs are also involved in chronic pathological processes observed in neurodegenerative diseases and neuroinflammation [15-17]. For example, the PVF activities are altered preceding the onset of amyotrophic lateral sclerosis [18]. In human Alzheimer’s disease, different locations of FBs show distinct perturbations of gene profiles, with leptomeningeal FBs enriched with solute transporters and PVF enriched with ECM expression [19].

Previous studies on cerebral, pulmonary, and cardiac FBs have shown that Ca^2+^ plays a critical role in various essential functions of FBs, including differentiation into myofibroblasts, proliferation, migration, fibrotic gene expression, and contraction [20-24]. Intracellular Ca^2+^ signals in FBs are generated by Ca^2+^ entry through channels in the plasma membrane and Ca^2+^ release from intracellular stores [22]. Studies have shown that Ca^2+^ influx mediated by transient receptor potential (TRP) channels plays an important role in FB differentiation [25]. Furthermore, disruption of Ca^2+^ signaling in FBs by nifedipine could attenuate pulmonary fibrosis [26]. Overall, proper regulation of intracellular Ca^2+^ levels is crucial for maintaining FB homeostasis and function.

Despite these aforementioned efforts, the detailed functional roles and the signaling mechanisms of different FB subgroups remain in the early stages of exploration. In this study, we utilized *in vivo* two-photon microscopy (TPM) in awake transgenic mice that express Ca^2+^ fluorescent indicators in three subgroups of leptomeningeal FBs: ArachF, PiaF, and PVF. We aimed to investigate their Ca^2+^ signaling characteristics under physiological conditions and during systematic inflammation to gain further insights into their potential underlying mechanism. Addressing these questions will significantly expand our understanding of normal CNS physiology and further elucidate pathological conditions such as inflammatory and degenerative diseases affecting the leptomeninges.

## Results

### Leptomeningeal fibroblast subgroups have distinct locations and ultrastructures

To investigate the three leptomeningeal FB subgroups ArachF, PiaF and PVF, we used a mouse line (*Cdh5*-GCaMP8) with a genetically encoded green Ca^2+^ sensitive protein (GCaMP8) under the Cadherine-5 promoter (*Cdh5*), a marker of adherens junctions. *Cdh5* is highly expressed in vascular endothelial cells, serves as a landmark for the leptomeninges, and transcriptomic studies have indicated a notable expression in brain FBs as well [5, 27, 28]. Our immunohistochemical staining of these mice showed co-localization of FB marker PDGFRα (cyan) and GCaMP8 expression (green) in the leptomeninges and as a thin layer outside vascular endothelial cells (red) (**Fig. 1A**). We performed *in vivo* TPM in the *Cdh5*-GCaMP8 mice with chronic cranial window and intravenous injection of Texas Red and used second-harmonic generation (SHG) to visualize the thick collagen layer in dura mater. The distinct locations of FBs in the leptomeninges allowed us to identify ArachF, PiaF, and PVF under TPM (**Fig. 1B x-z plane** and **Sup. Fig. 1**B **x-y plane**). ArachFs form the arachnoid mater, the outer layer above the pial vessels. PiaFs form the pia mater beneath the pial vessels. However, in areas lacking pial vessels, PiaFs and ArachFs are positioned closer together. PVFs envelop pial vessels but are mostly separated from the endothelial layers by a layer of vascular mural cells (**Fig. 1B**).

**Figure 1.**
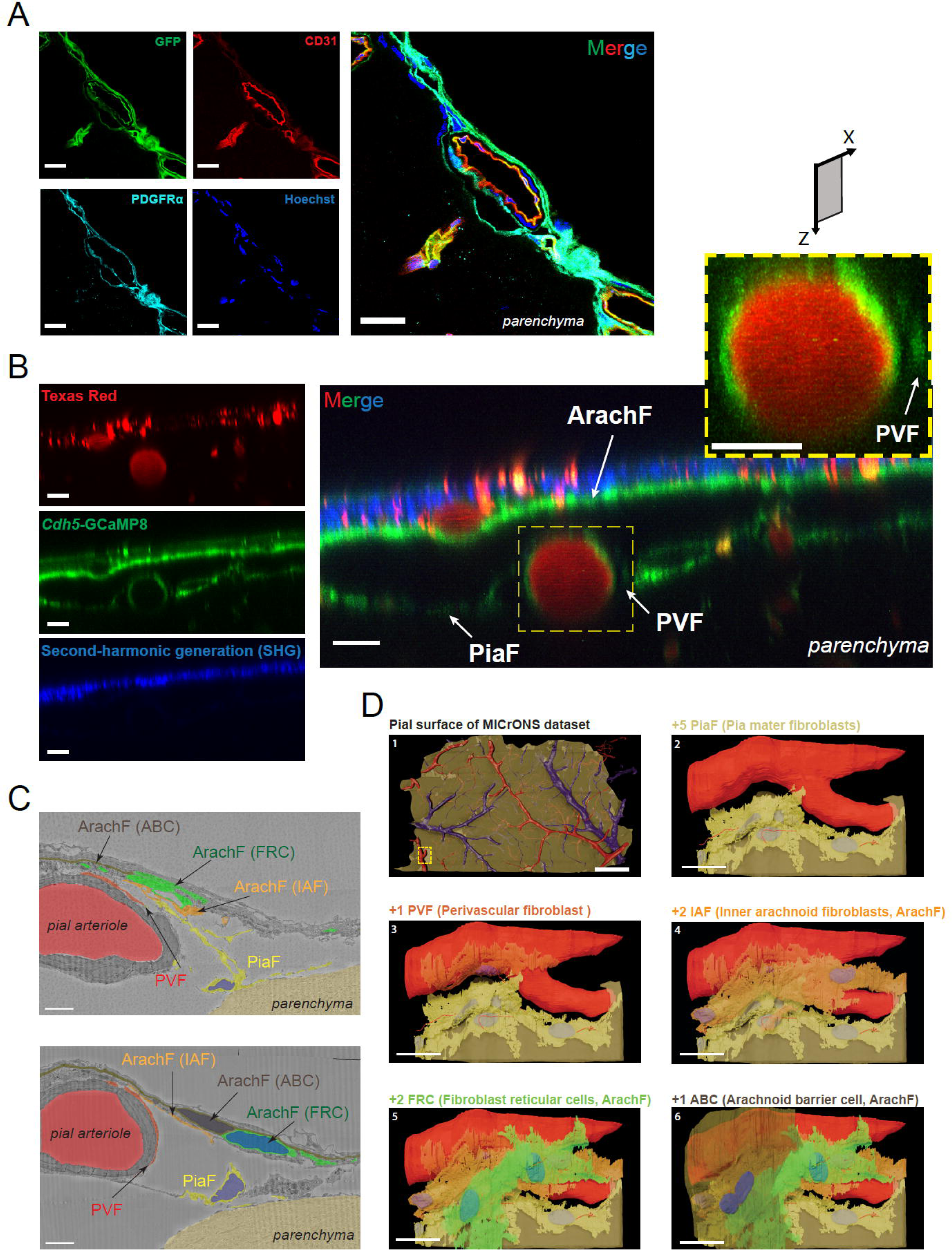
Anatomical classification of leptomeningeal fibroblast subgroups. (**A**) Immunohistochemical staining of brain slices of mouse *Cdh5*-GCaMP8. Green: *Cdh5*-GCaMP8. Red: CD31 for endothelial cells. Cyan: PDGFRα. Blue: Hoechst. Scale bar: 20 μm. (**B**) TPM of leptomeningeal fibroblasts (FBs) across the x-z plane. Red: i.v. injection of Texas Red for vessel lumen. Green: *Cdh5*-GCaMP8. Blue: second-harmonic generation. Based on the FB location in TPM images, the meningeal FBs are classified as arachnoid FB (ArachF), pia mater FB (PiaF), and perivascular FB (PVF). Scale bar: 20 μm. (**C**) MICrONS database of high-resolution electron microscopy (EM) shows clear ultrastructure of leptomeningeal FBs. The arachnoid mater separates the subarachnoid space from the subdural space, and contains FB reticular cells (FRC), inner arachnoid FBs (IAF) and arachnoid barrier cells (ABC). FRC, IAF, and ABC are subtypes of ArachF. As it is impossible to separate them by TPM, we only distinguish them in the EM dataset. Scale bar: 5 μm. (**D**) Sequential addition of different structures in the EM dataset (snapshots of Sup. Video 1). Top left: large-scale pial surface. Scale bar: 200 μm. Top right: Addition of PiaF. Middle left: further addition of PVF. Middle right: further addition of IAF (a subtype of ArachF). Bottom left: further addition of FRC (a subtype of ArachF). Bottom right: further addition of ABC (a subtype of ArachF). Scale bar: 20 μm.

We investigated the ultrastructures and 3D structures of ArachF, PiaF and PVF using the MICrONS dataset of large-scale high-resolution electron microscopy (**Fig. 1C, Sup. Video. 1 and Sub Fig. 2**) [29]. We selected a region containing a relatively large pial arteriole branching into a penetrating arteriole (**Fig. 1D**). We segmented PVF and PiaF based on whether they encircled the vasculature (PVF) or covered the pial surface (PiaF). We identified three subgroups of ArachF based on their locations and ultrastructural characteristics in the arachnoid mater: Arachnoid barrier cell (ABC), FB reticular cells (FRC), and inner arachnoid FBs (IAF) [2-5]. Based on the sequential 2D electron microscopy images of this leptomeningeal region, we made a 3D reconstruction of those FBs, identifying their association with the pia mater, a pial arteriole, or the arachnoid mater (**Fig. 1C-D** and **Sup. Video. 2**). These ultrastructural analyses demonstrate that the different subgroups of leptomeningeal FBs, ArachF (ABC, IAF, FRC), PVF and PiaF, have specific locations and/or unique ultrastructural features, suggesting that they may serve distinct functional roles in both physiological and pathological conditions (**Sup. Fig. 1**A).

**Figure 2.**
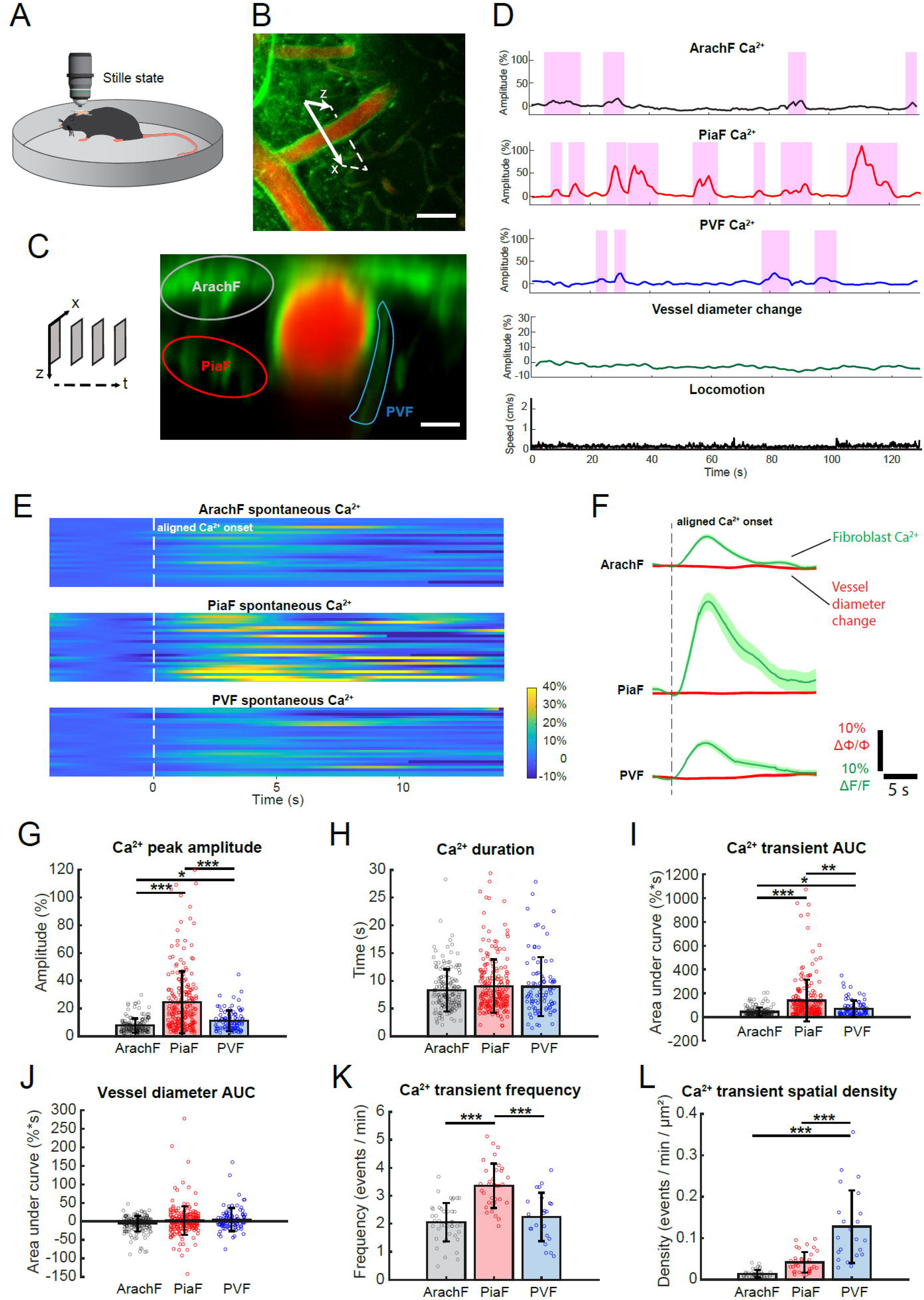
Distinct profiles of spontaneous Ca^2+^ transients for each fibroblast subgroup. (**A**) Diagram of an awake mouse implanted with a chronic cranial window and imaged by TPM. The spontaneous Ca^2+^ transients were recorded when the mouse was at a stilled (resting) state without locomotion. (**B**) Representative TPM image at the x-y plane. To simultaneously record Ca^2+^ activities across all three fibroblast (FB) subgroups, along with vessel diameter changes, we performed time-series x-z plane imaging. Scale bar: 50 μm. (**C**) Representative TPM image at x-z plane, with marked FB subgroups: ArachF, PiaF, and PVF. Scale bar: 10 μm. (**D**) Representative traces of Ca^2+^ activities at ArachF, PiaF, and PVF are marked in (C), along with co-localized pial vessel diameter and synchronized locomotion recording. (**E**) Heat maps of spontaneous Ca^2+^ transients at each FB subgroup, by aligning Ca^2+^ onsets. X-axis: time. Y-axis: trace number. (**F**) Overlaid Ca^2+^ and vessel diameter traces for each FB subgroups. The shadow represents s.e.m. of all traces. (**G-L**) Comparison of spontaneous Ca^2+^ profiles at each FB subgroup, on (**G**) Ca^2+^ peak amplitude, (**H**) Ca^2+^ transient area under the curve (AUC), (**I**) Ca^2+^ transient duration, (**J**) Vessel diameter change, (**K**) Ca^2+^ transient frequency, and (**L**) Ca^2+^ transient spatial density. (ArachF: N = 7 mice, n = 42 locations, n′ = 174 events; PiaF: N = 7 mice, n = 34 locations, n′ = 215 events; PVF: N = 7 mice, n = 22 locations, n′ = 91 events.) Bar plots are presented as the mean ± s.t.d. Linear mixed-effect models were used, followed by Tukey post hoc tests for pairwise comparisons. * p < 0.05, ** p < 0.01, *** p < 0.001.

### Different Ca^2+^ profiles of fibroblast subgroups at the resting state

To investigate whether the three subgroups of FBs have different Ca^2+^ signaling features, we recorded Ca^2+^ signals by acquiring sequential x-z plane TPM images of awake mice with chronic cranial windows, while simultaneously monitoring their locomotion. We analyzed Ca^2+^ signals of ArachF, PiaF and PVF at the resting state, i.e., the mice remained still (**Fig. 2A-C** and **Sup. Video. 3**). Representative traces of Ca^2+^ signals at each leptomeningeal FB subgroup, along with synchronized recording of co-localized vessel diameter and locomotion, are shown in **Fig. 2D**. These traces indicated the occurrence of spontaneous Ca^2+^ transients in the absence of active mouse behaviors and vessel diameter changes. Next, we aligned the onsets of Ca^2+^ transients for each FB subgroup (**Fig. 2E**) and averaged Ca^2+^ transients with synchronized vessel diameter changes (**Fig. 2F**). Ca^2+^ transient peak amplitudes and area under the curves (AUCs) were the largest for PiaF among all subgroups, and those for PVF were significantly larger than ArachF (**Fig. 2G, I**). Interestingly, the duration of spontaneous Ca^2+^ transients across all subgroups was similar, although it ranged from 1-2 seconds to as long as 30 seconds (**Fig. 2H**). In contrast, the AUCs of vessel diameter changes remained close to zero (**Fig. 2J**). Further, we studied Ca^2+^ transient frequency, i.e., the number of transients per minute, which showed significantly more frequent Ca^2+^ activities at PiaF compared with the other two subgroups (**Fig. 2K**). Considering that the surface area of ArachF, PiaF and PVF differed greatly at x-z plane, we estimated the surface area of each FB subgroup in each recording and normalized the Ca^2+^ frequencies by area to obtain spatial density of Ca^2+^ transients. PVF Ca^2+^ showed the largest spatial density compared with the other two subgroups (**Fig. 2L**).

Next, we examined whether spontaneous Ca^2+^ signals are synchronized within the same FB subgroup or across different subgroups in neighboring locations. For this purpose, we placed regions of interest (ROIs) over each FB subgroup in the same TPM recording (**Sub. Fig. 3A**) and plotted the Ca^2+^ traces of each ROI (**Sub. Fig. 3B**). We calculated the correlation coefficient for every pair of ROIs and plotted a heat map in two different modes: sorted by FB subgroups (**Sub. Fig. 3C**) and sorted by the spatial proximity of the ROIs (**Sub. Fig. 3D**). Interestingly, both heat maps lacked clusters of highly correlated ROIs, indicating that FB Ca^2+^ activities are generated independently, regardless of spatial proximity or subgroup. In addition to this representative case, we also calculated the cross-correlations of all Ca^2+^ traces between each pair of subgroups within the same TPM recordings (**Sub. Fig. 4**). Similarly, we showed random and dispersed cross-correlation peaks across the recording locations, further supporting the idea that FB Ca^2+^ activity occurs independently across space and subgroups.

**Figure 3.**
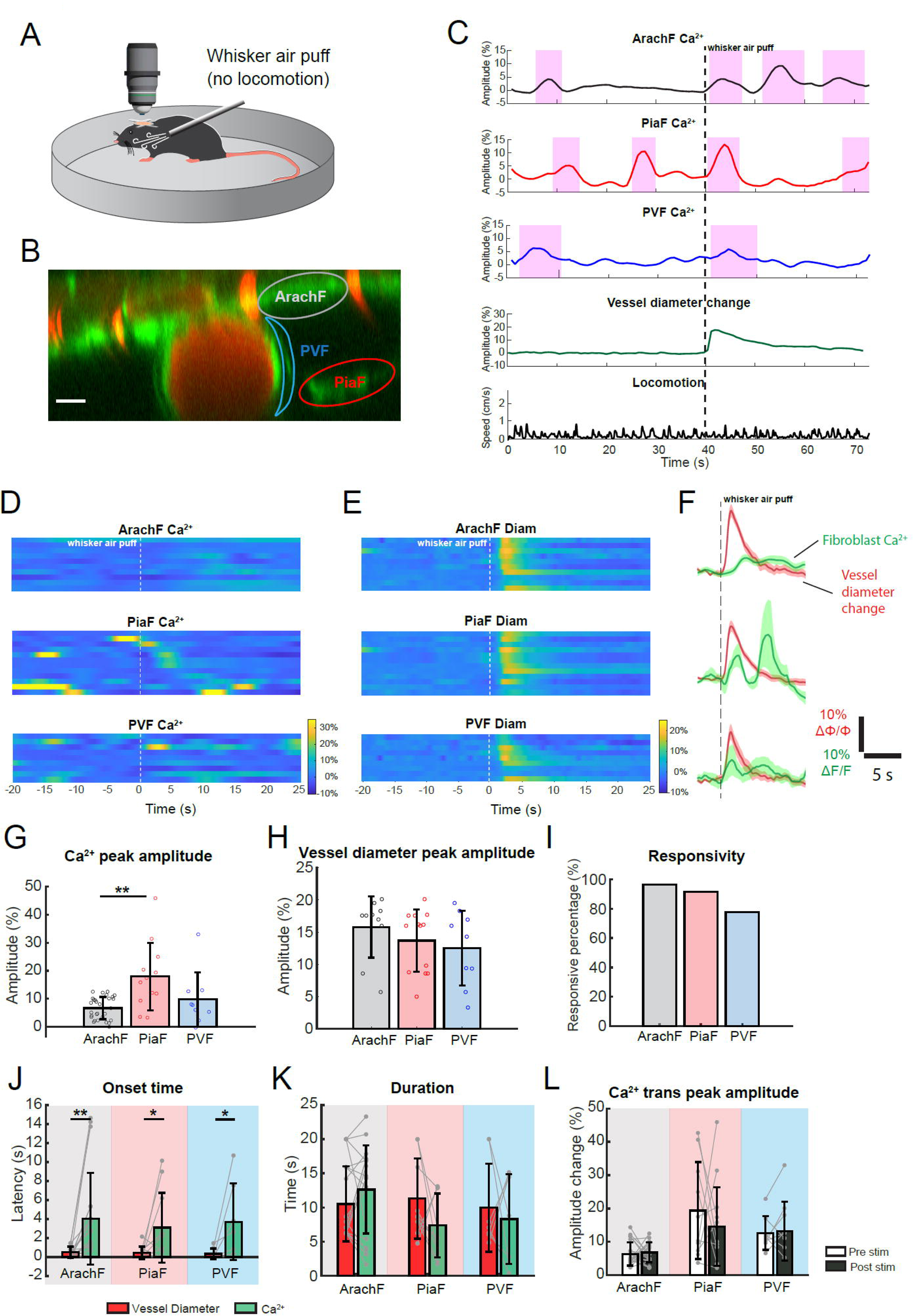
Whisker air puff elicits pial arteriole dilation and fibroblast Ca^2+^ activities. (**A**) Diagram of an awake mouse stimulated by whisker air puff during TPM. (**B**) A representative TPM image with marked fibroblast (FB) subgroups. Scale bar: 10 μm. (**C**) Representative traces of Ca^2+^ activities at ArachF, PiaF, and PVF are marked in (B), along with co-localized pial vessel diameter and synchronized locomotion recording. (**D-E**) Heat maps of air-puff stimulation-elicited Ca^2+^ transients (D) and vessel diameter change (E) at each FB subgroup, by aligning air puff onset. x-axis: time. Y-axis: trace number. (**F**) Overlaid Ca^2+^ and vessel diameter traces for each FB subgroups. The shadow represents s.e.m. of all traces. (**G-I**) Comparison of air-puff induced Ca^2+^ responses at each FB subgroup, on (**G**) Ca^2+^ peak amplitude, (**H**) Vessel diameter change and (**I**) Ca^2+^ responsivity. (**J-K**) Comparison between vessel diameter change and air-puff induced FB Ca^2+^, on onset time (J) and duration (K). (**L**) Comparison of FB Ca^2+^ transient peak amplitudes before and after air-puff stimulation. (ArachF: N = 4 mice, n = 9 locations, n′ = 29 trials; PiaF: N = 3 mice, n = 8 locations, n′ = 13 trials; PVF: N = 4 mice, n = 7 locations, n′ = 8 trials.) Bar plots are presented as the mean ± s.t.d. Linear mixed-effect models were used followed by Tukey post hoc tests for pairwise comparisons. * p < 0.05, ** p < 0.01.

**Figure 4.**
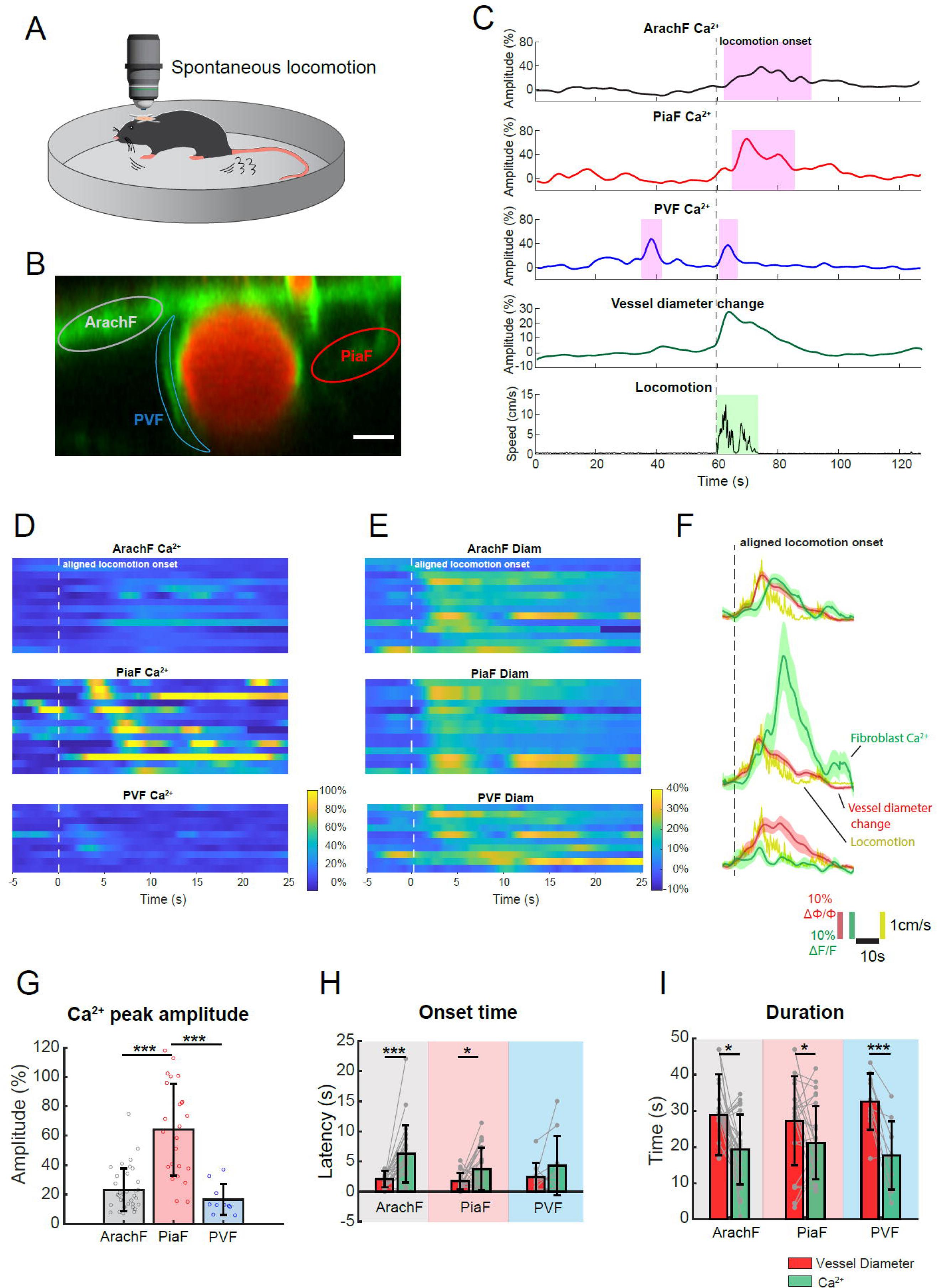
Spontaneous locomotion elicits strong vasodilation and fibroblast Ca^2+^ activities. (**A**) Diagram of an awake mouse with spontaneous locomotion during TPM. (**B**) A representative TPM image with marked fibroblast (FB) subgroups. Scale bar: 10 μm. (**C**) Representative traces of Ca^2+^ activities at ArachF, PiaF, and PVF are marked in (B), along with co-localized pial vessel diameter and synchronized locomotion recording. (**D-E**) Heat maps of locomotion-elicited Ca^2+^ transients (D) and vessel diameter change (E) at each FB subgroup, by aligning locomotion onset. x-axis: time. Y-axis: trace number. (**F**) Overlaid Ca^2+^, vessel diameter, and locomotion traces for each FB subgroups. The shadow represents s.e.m. of all traces. (**G**) Comparison of locomotion-induced Ca^2+^ responses at each FB subgroup, focusing on Ca^2+^ peak amplitude. (**H-I**) Comparison between vessel diameter change and locomotion-induced FB Ca^2+^, on onset time (H) and duration (I). (ArachF: N = 5 mice, n = 7 locations, n′ = 35 trials; PiaF: N = 6 mice, n = 9 locations, n′ = 25 trials; PVF: N = 4 mice, n = 8 locations, n′ = 10 trials.) Bar plots are presented as the mean ± s.t.d. Linear mixed-effect models were used, followed by Tukey post hoc tests for pairwise comparisons. * p < 0.05, ** p < 0.01, *** p < 0.001.

### Whisker air puff induced pial vessel dilation and fibroblast Ca^2+^ rise

Whisker air puff is widely used to induce neurovascular coupling in the somatosensory cortex of awake mice. We asked whether FB Ca^2+^ is involved in this process and whether different FB subgroups exhibit distinct responses. After habituating the awake mice to a modest (∼4 psi) and short (1 s) whisker air puff, we recorded TPM x-z plane images and analyzed trials in which the mouse whiskers responded to an air puff without accompanying locomotion (**Fig. 3A**). Whisker air puff stimulation reliably induced pial arteriole dilation, accompanied by Ca^2+^ transients in all three FB subgroups (**Fig. 3B-C**). Collectively, we visualized and averaged the Ca^2+^ responses and their co-localized arteriole diameter changes for each FB subgroup (**Fig. 3D-F**). The peak amplitudes of Ca^2+^ responses of PiaF were significantly higher than those of ArachF (**Fig. 3G**), while the vessel diameter changes were similar across the three cases (**Fig. 3H**). Interestingly, although PVFs are closest to the pial arteries, their responsivity, defined as the proportion of triggered Ca^2+^ transients across all trials, was lowest among the three subgroups (**Fig. 3I**). Next, we investigated the temporal relationship between vessel diameter changes and associated FB Ca^2+^ activity. The onset of triggered Ca^2+^ transients was significantly delayed compared to the onset of vasodilation across all FB subgroups (**Fig. 3J**), while their response durations were similar (**Fig. 3K**). Lastly, we examined whether the whisker air puff-triggered Ca^2+^ kinetics differ from spontaneous Ca^2+^ events, by comparing Ca^2+^ peak amplitudes (**Fig. 3L**), Ca^2+^ transient durations (**Sub. Fig. 5A**), and Ca^2+^ transient AUCs (**Sub. Fig. 5B**) before and after air puff stimulation. None of these comparisons showed statistically significant differences. Overall, these results suggested that whisker air puffs induce preceding pial arteriole dilation and followed by FB Ca^2+^ activations across all leptomeningeal FB subgroups. The characteristics of the induced FB Ca^2+^ events are similar to the spontaneous Ca^2+^ events in FBs.

**Figure 5.**
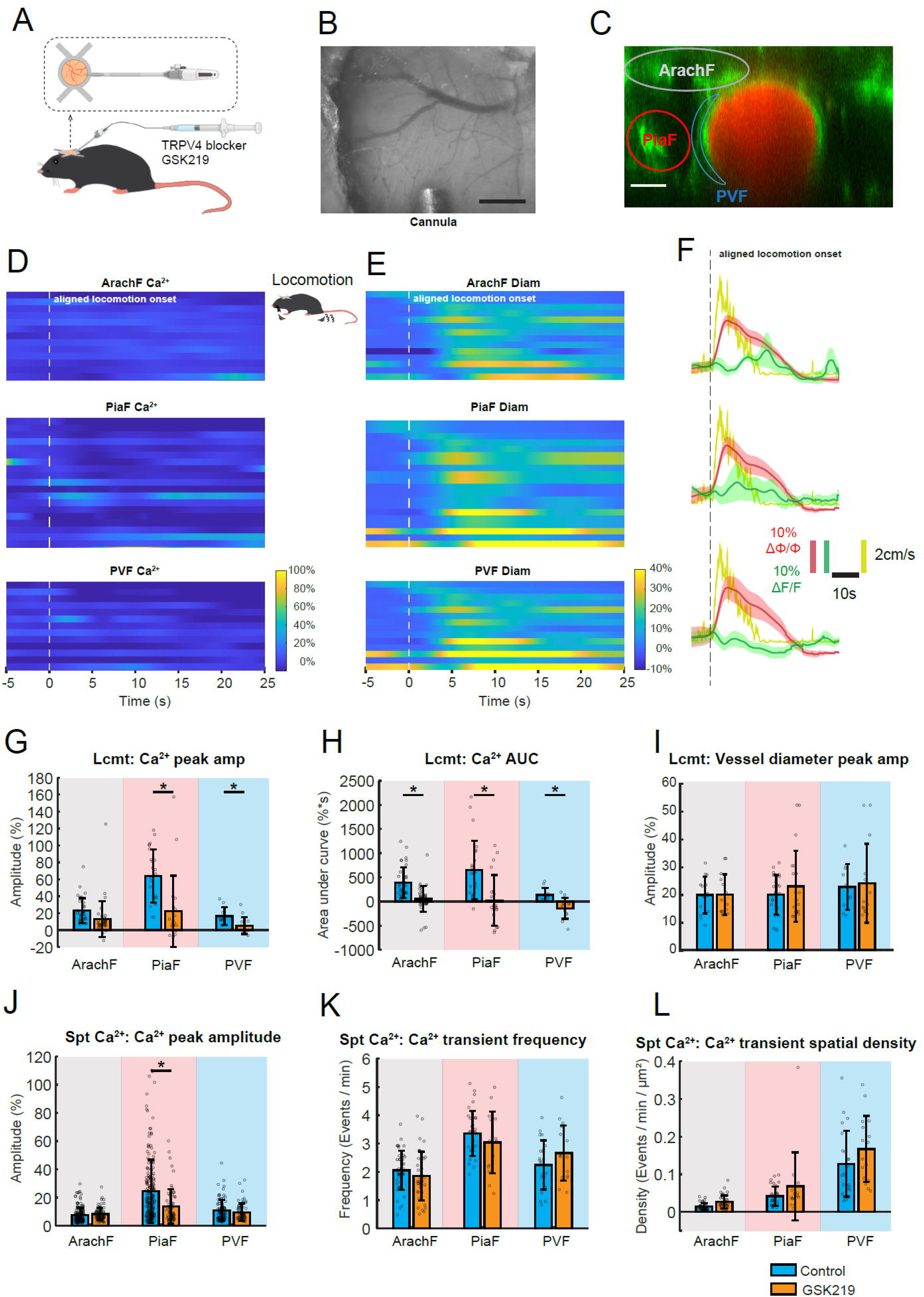
TRPV4 channels mediate locomotion-induced fibroblast Ca^2+^ activities. (**A**) Diagram of an awake mouse chronically implanted with a cannula. (**B**) Wide-field imaging of an implanted cannula beneath a chronic cranial glass window. Scale bar: 500 μm. (**C**) A representative TPM image with marked fibroblast (FB) subgroups after TRPV4. Scale bar: 15 μm. (**D-E**) After administration of GSK219, heat maps of locomotion-elicited Ca^2+^ transients (D) and vessel diameter change (E) at each FB subgroup, by aligning locomotion onset. x-axis: time. Y-axis: trace number. (**F**) Overlaid Ca^2+^, vessel diameter, and locomotion traces for each FB subgroups after administration of GSK219. The shadow represents s.e.m. of all traces. (**G-I**) Comparison of locomotion-induced Ca^2+^ responses without or with intracortical infusion of GSK219, focusing on (G) Ca^2+^ peak amplitude, (H) Ca^2+^ transient AUC, and (I) Peak amplitude of vessel diameter change. (**J-L**) Comparison of spontaneous Ca^2+^ transients without or with intracortical infusion of GSK219, focusing on (J) Ca^2+^ peak amplitude, (K) Ca^2+^ transient frequency, and (L) Ca^2+^ transient spatial density. (**GSK219 spontaneous Ca^2+^** - ArachF: N = 5 mice, n = 11 locations, n′ = 93 events; PiaF: N = 5 mice, n = 11 locations, n′ = 63 events; PVF: N = 5 mice, n = 11 locations, n′ = 61 events.) (**GSK219 locomotion -** ArachF: N = 4 mice, n = 11 locations, n′ = 38 trials; PiaF: N = 3 mice, n = 9 locations, n′ = 21 trials; PVF: N = 3 mice, n = 10 locations, n′ = 14 trials.) Bar plots are presented as the mean ± s.t.d. Linear mixed-effect models were used, followed by Tukey post hoc tests for pairwise comparisons. * p < 0.05, ** p < 0.01, *** p < 0.001. The diagram of the cannula is generated using BioRender.

### Spontaneous locomotion elicited strong vasodilation and fibroblast Ca^2+^ activities

Spontaneous locomotion frequently occurs in awake mice and is always accompanied by volitional whisking [30]. Unlike artificial whisker air puff, spontaneous locomotion represents a more natural behavior. Therefore, we asked how each FB subgroup responds to spontaneous locomotion. For this purpose, we continuously performed TPM and captured spontaneous locomotion of awake mice (**Fig. 4A**). Similar to whisker air puff stimulation, spontaneous locomotion induced neurovascular coupling leads to pial arteriole dilation and Ca^2+^ elevations across all FB subgroups (**Fig. 4B-C** and **Sup. Video. 4**). By aligning locomotion onsets, we overlaid FB Ca^2+^ traces, vessel diameter changes and locomotion, revealing a strong and time-locked response to locomotion (**Fig. 4D-F**). Further quantification showed that Ca^2+^ peak amplitudes were highest in PiaF (**Fig. 4G**), although arteriole diameter changes were similar across FB subgroups (**Sub. Fig. 6A**). The onset of vasodilation preceded that of elicited Ca^2+^ transients for both ArachF and PiaF (**Fig. 4H**). Moreover, the duration of vasodilation was significantly longer than that of elicited Ca^2+^ activities across all FB subgroups (**Fig. 4I**). Considering that a single locomotion event can trigger single or multiple Ca^2+^ transients, we asked whether the occurrence of FB Ca^2+^ transients predominantly coincide with each phase of the vasodilation — namely, onset, peak and offset. Thereby, we digitized the onset of each Ca^2+^ transient as a vertical line, and overlaid all the transient onsets aligned to the onset, offset and peak time of vessel diameter changes (**Sub. Fig. 7A-I**), and further summarized them as bar plots (**Sub. Fig. 7J-R**). These data clearly indicated that the occurrence of FB Ca^2+^ transients at all three subgroups are correlated within 0 –5 seconds following vasodilation onset, but not with the offset or peak time. Lastly, we compared the three scenarios: spontaneous Ca^2+^, whisker air puff and locomotion (**Sub. Fig. 6B-C**). Locomotion elicited the strongest vessel dilation, greater than whisker air puff stimulation; while spontaneous Ca^2+^ events were not accompanied with vessel diameter changes (**Sub. Fig. 6B**). In contrast, Ca^2+^ transient amplitudes were highest during locomotion for ArachF and PiaF, while those for PVF remained consistent across three scenarios (**Sub. Fig. 6C**). Together, these results suggest that spontaneous locomotion induces strong pial arteriole dilation and FB Ca^2+^ transients. Vasodilation onset precedes FB Ca^2+^ activity, mostly within a 5-second time window, and Ca^2+^ transients at PiaF consistently exhibit the strongest responses.

**Figure 6.**
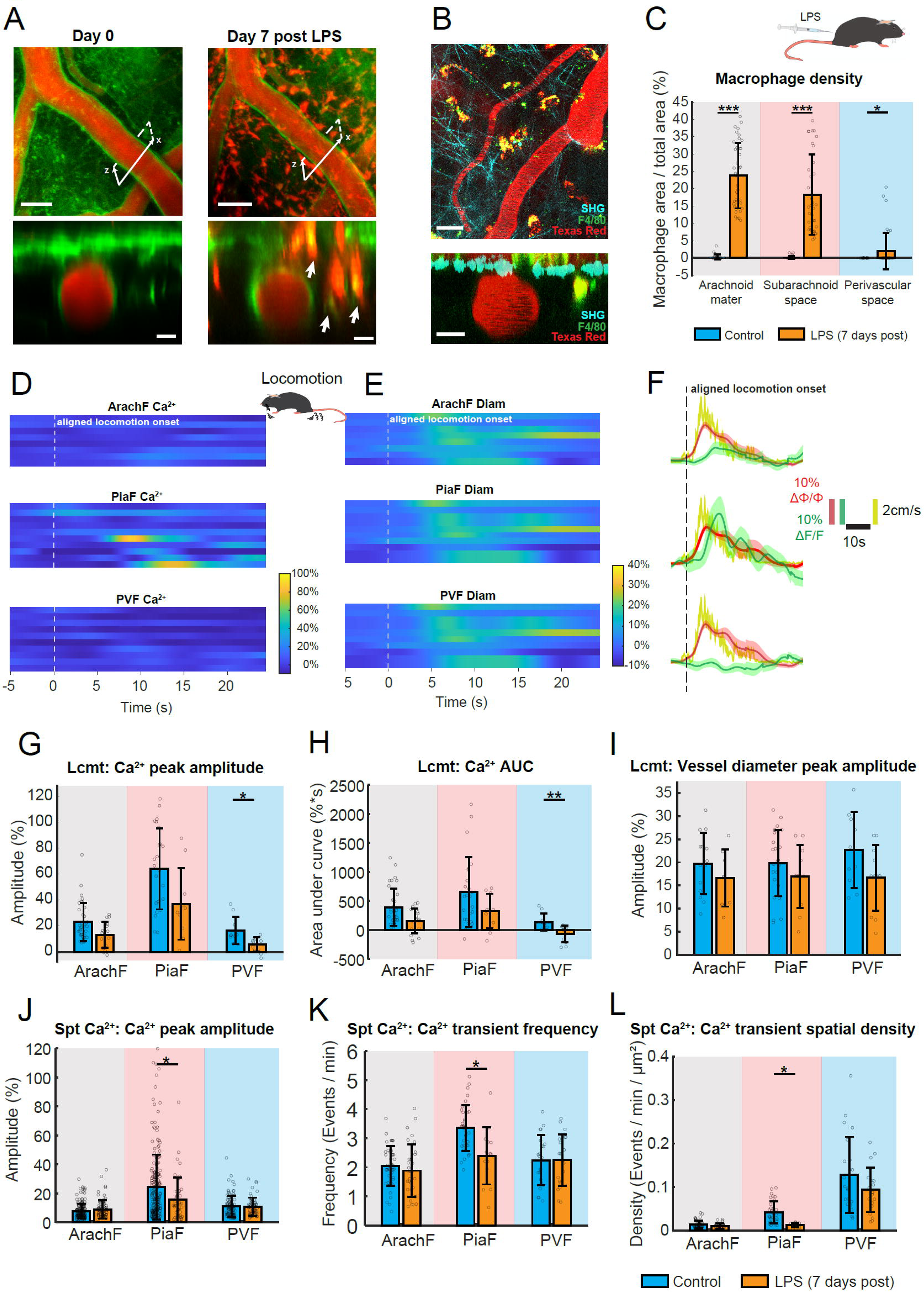
Systematic inflammation alters Ca^2+^ activities in different fibroblast subgroups. (**A**) Comparison of brain surfaces of mice before and 7 days after LPS administration. Top: large-scale x-y plane image (Scale bar: 50 μm). Bottom: small-scale x-z plane image (Scale bar: 10 μm). (**B**) Identification of leptomeningeal macrophages that engulf red fluorescent dye leaked from the disrupted blood-brain barrier. SHG: second harmonic generation. F4/80 or F4/80-Alexa488: macrophage. Texas Red: vessel lumen dye. Scale bar: Top 25 μm; bottom: 10 μm. (**C**) Top: Diagram of an awake mouse with intraperitoneal injection of LPS to induce systemic inflammation. Bottom: macrophage density at arachnoid mater, subarachnoid space, and perivascular space before and 7 days after administration of LPS. (**D-E**) After administration of LPS, heat maps of locomotion-elicited Ca^2+^ transients (D) and vessel diameter change (E) at each FB subgroup, by aligning locomotion onset. X-axis: time. Y-axis: trace number. (**F**) Overlaid Ca^2+^, vessel diameter, and locomotion traces for each FB subgroups after administration of LPS. The shadow represents s.e.m. of all traces. (**G-I**) Comparison of locomotion-induced Ca^2+^ responses before and 1 week after LPS, focusing on (G) Ca^2+^ peak amplitude, (H) Ca^2+^ transient AUC, and (I) Peak amplitude of vessel diameter change. (**J-L**) Comparison of spontaneous Ca^2+^ transients before and 1 week after LPS, focusing on (J) Ca^2+^ peak amplitude, (K) Ca^2+^ transient frequency, and (L) Ca^2+^ transient spatial density. (**LPS spontaneous Ca^2+^** -ArachF: N = 3 mice, n = 9 locations, n′ = 80 events; PiaF: N = 3 mice, n = 8 locations, n′ = 46 events; PVF: N = 3 mice, n = 11 locations, n′ = 62 events.) (**LPS locomotion -** ArachF: N = 3 mice, n = 7 locations, n′ = 18 trials; PiaF: N = 3 mice, n = 7 locations, n′ = 11 trials; PVF: N = 3 mice, n = 8 locations, n′ = 11 trials.) Bar plots are presented as the mean ± s.t.d. Linear mixed-effect models were used, followed by Tukey post hoc tests for pairwise comparisons. * p < 0.05, ** p < 0.01, *** p < 0.001.

### TRPV4 channels mediate locomotion-induced fibroblast Ca^2+^ activity

We further investigated the mechanisms underlying locomotion-induced FB Ca^2+^ activity. Previous studies have shown that TRPV4 channels are expressed in FBs in different organs and are involved in the function of mechanosensors [25, 31]. Therefore, we hypothesized that TRPV4 channels may also play a role in leptomeninges in this process. To test this, we developed a cannula delivery system, by which a cannula was chronically implanted through the cover glass and inserted into the parenchyma, allowing TPM recording while administering the drug (**Fig. 5A-B**). Twenty minutes after the slow infusion of the TRPV4 antagonist GSK219 (GSK2193874) through the cannula delivery system, we examined FB Ca^2+^ activities during locomotion (**Fig. 5C-F**). Locomotion-induced FB Ca^2+^ responses were significantly attenuated by GSK219 across all three FB subgroups (**Fig. 5G-H**), whereas locomotion-induced pial arteriole dilation remained unchanged between control and GSK219-treated conditions (**Fig. 5I, Sub. Fig. 8C**).

Next, we asked whether GSK219 alters spontaneous FB Ca^2+^ activity. After sorting and aligning all spontaneous Ca^2+^ and vessel diameter traces by Ca^2+^ onset (**Sub. Fig. 8A-B**), we observed a significant reduction in Ca^2+^ peak amplitudes and AUCs at PiaF following GSK219 treatment (**Fig. 5J, Sub. Fig. 8E**), but not at ArachF or PVF. Vessel diameters remained unchanged in all cases (**Sub. Fig. 8D**). Interestingly, further comparisons of spontaneous Ca^2+^ transient frequency (**Fig. 5K**), Ca^2+^ transient spatial density (**Fig. 5L**) and Ca^2+^ active period (**Sub. Fig. 8F)**, defined as the percentage of Ca^2+^-active duration relative to the total TPM recording time, showed no significant change by GSK219.

These results suggest that TRPV4 channels mediate locomotion-induced FB Ca^2+^ activity. They contribute to enhancing spontaneous Ca^2+^ transient amplitudes at PiaF but are largely uninvolved in the generation of spontaneous Ca^2+^ events across all FB subgroups.

### Systematic inflammation alters fibroblast Ca^2+^ activity

Growing evidence has indicated important roles for FBs in brain injury and neuroinflammation [15]. We investigated whether FB Ca^2+^ activity is involved in brain immune response. To induce systemic inflammation, we administered lipopolysaccharide (LPS) via intraperitoneal (i.p.) injection in mice with chronic cranial windows. Six hours after LPS injection, TPM revealed a severe blood–brain barrier leakage (**Sub. Fig. 9A**), which significantly reduced the two-photon laser penetration into brain tissue, thereby preventing us from performing x-z TPM imaging. The disrupted blood-brain barrier (BBB) persisted for several days. Therefore, we recorded FB Ca^2+^ activities on day 7 post-LPS administration (**Fig. 6A**).

Compared with pre-LPS, the cortical surface of mice 7 days post-LPS is still covered by a large number of red fluorescent cells, and they are located above, within, and beneath the arachnoid layer (**Fig. 6A**, bottom right). As the key immune function of macrophages is phagocytosis [32, 33], macrophages likely engulf the red fluorescent dye in the parenchyma, which is leaked from the disrupted BBB. To further examine this, we intravenously injected macrophage antibody F4/80-Alexa488 in mice 12 hours before the acute *in vivo* TPM experiment (see detailed protocol in the Methods section). We observed highly co-localized macrophages (green) and Texas Red (**Fig. 6B**), which is not due to fluorescence bleed-through (**Sub. Fig. 9B**). This suggests that the ‘red cells’ in Fig.6A were macrophages. In particular, some of those macrophages contact closely with ArachF and PiaF (**Fig. 6A**, white arrows), or in the process of infiltration across the dura and potential the arachnoid mater (**Sup. Video. 5**). High-resolution 2D and 3D ultrastructures from the MICrONS electron microscopy database also revealed that macrophages are embedded within the subarachnoid space, in close contact with leptomeningeal fibroblasts (**Sup. Fig. 10**). Next, we quantified the density of Texas Red-positive macrophages in each FB region by calculating the ratio of macrophage area to the total area in x-z TPM images. Our results showed a markedly elevated macrophage density within all the leptomeningeal sublayers 7 days after LPS administration (**Fig. 6C**).

Furthermore, we examined locomotion-induced FB Ca^2+^ activities at day 7 (**Fig. 6D-F**). The peak amplitudes and AUCs of Ca^2+^ transients elicited by locomotion showed a decreasing trend across all three FB subgroups compared to controls, but reached statistical significance only for PVF (**Fig. 6G-H**). In comparison, vasodilation remained consistent in peak amplitudes and AUCs between control and LPS-treated mice (**Fig. 6I, Sub. Fig. 11C**). Next, we asked whether systematic inflammation altered the spontaneous Ca^2+^ profiles of the FBs. The overlay of spontaneous FB Ca^2+^ transients by their onsets (**Sub. Fig. 11A-B**) and further quantification showed a significant change in PiaF Ca^2+^ transients, but not ArachF or PVF. Spontaneous Ca^2+^ peak amplitude (**Fig. 6J**), transient frequency (**Fig. 6K**), transient spatial density (**Fig. 6L**) and Ca^2+^ active period (**Sub. Fig. 11E**) at PiaF were significantly decreased. Overall, systematic inflammation by LPS induces a decreased spontaneous Ca^2+^ activity of PiaF but not ArachF and PVF, while it affects locomotion-induced Ca^2+^ amplitudes of PVF but not ArachF and PiaF.

## Discussion

In this study, we identified leptomeningeal FBs and investigated their spontaneous Ca^2+^ transient profiles and behavior-elicited responses in awake mice. We separated leptomeningeal FBs into three subgroups - ArachF, PiaF and PVF, based on their anatomically distinct locations revealed by electron microscopy and TPM, with ArachF containing 3 different subgroups of FBs. We found that PiaF had the highest spontaneous and behavior-elicited Ca^2+^ activity amplitude among the three subgroups.

Furthermore, a TRPV4 blocker inhibited locomotion-induced FB Ca^2+^ transients, initiating seconds after pial arteriole vasodilation. Finally, we found that systematic inflammation altered both spontaneous and locomotion-induced Ca^2+^ activity of PVF and PiaF, which is likely attributed to the infiltration of macrophages. Our results provide important and novel insight into the Ca^2+^ signaling and their possible function of leptomeningeal FBs.

### Spontaneous Ca^2+^ activity of leptomeningeal fibroblasts at rest

In our TPM experiments, we employed a novel scanning mode —x-z scanning over time, which enabled us to observe the calcium dynamics of leptomeningeal FB subgroups with relatively high temporal resolution in awake *Cdh5*-GCaMP8 mouse brains. At rest, we found that the spontaneous FB Ca^2+^ profiles differed among the three subgroups. Although their Ca^2+^ transients exhibited similar durations, other parameters, like peak amplitude, temporal frequency, and spatial density, revealed subgroup-specific differences.

The similarity in Ca^2+^ transient duration implies a conserved mechanism governing onset and offset of these events across subgroups, suggesting that they might share closely related functional roles, such as extracellular matrix formation, tissue repair, and inflammation modulation [34-36]. Furthermore, we did not observe any synchronization of spontaneous Ca^2+^ activities in adjacent regions, either within or across different subgroups (**Sup. Fig. 3** and 4), providing general evidence that these cells lack intercellular gap junctions. This finding contrasts with a previous study reporting that inner arachnoid FBs are interconnected, and also connected with outer-layer cells, via gap junctions that facilitate ion exchange [2, 37, 38].

On the other hand, the observed differences in their Ca^2+^ profiles underscore the distinct identities of these FB subgroups. Each subgroup has unique compositions, locations, and inputs. The arachnoid mater consists of multiple FB layers, and the fact that the ArachFs lack innervation [38] and direct connection with glial limitans may account for their lower Ca^2+^ event frequency compared with PiaF.

Conversely, the PVFs are directly opposed to the vessel wall [2], raising the possibility that PVFs’ activity could be influenced by adjacent vascular smooth muscle cells, endothelial cells, perivascular macrophages, or astrocytic endfeet [24]. This could explain the highest spatial density of Ca^2+^ transients at PVFs (**Fig. 2L**). However, we can’t exclude the possibility that the drastic differences of cross-sectional area of each FB subgroup in x-z plane lead to the discrepancy, and further 3D estimation of surface area of each FB subgroup is needed. Finally, PiaFs exhibit the highest Ca²⁺ amplitude and a moderate spatiotemporal frequency, likely attributable to their proximity to the brain parenchyma, absence of gap junctions, and larger cellular volume compared to PVFs.

### Pial arteriole dilation possibly induces the Ca^2+^ rise in leptomeningeal fibroblasts

The delayed Ca^2+^ elevation relative to vessel dilation, along with its attenuation by blockage of mechanosensitive and osmosensitive TRPV4 channels, suggests that the Ca^2+^ transients likely arise from the mechanical stretch by vasodilation upon natural behavior. Consistent with spontaneous Ca^2+^ profiles, PiaFs exhibit the highest Ca^2+^ amplitude among all subgroups by both whisker air puff stimulation and spontaneous locomotion. This is reflected by a recent transcriptomic study [2], showing the highest RNA expression of internal Ca^2+^ store gene markers at BFB1-positive FBs (PiaFs and PVFs), e.g., *Itpr1* and *Itpr2* genes for internal Ca^2+^ store, *Ryr2* gene for Ca^2+^-induced Ca^2+^ release.

Notably, PiaFs and PVFs are genetically indistinguishable, yet PiaF Ca^2+^ responses are more pronounced. This difference may stem from variations in translational regulation of Ca^2+^-related channels and the relatively larger extracellular space surrounding PiaFs, which could facilitate more efficient Ca^2+^ influx, internal release, and recycling. Furthermore, spontaneous locomotion—a more potent stimulus for pial arteriole dilation than whisker air-puff stimulation—elicits markedly higher Ca²⁺ responses, indicating that Ca²⁺ elevation scales with the degree of mechanical stretch.

As FB Ca²⁺ signaling is associated with the synthesis of collagen, laminin, and other ECM components [20, 34, 39], they are critical for maintaining leptomeningeal structure. We hypothesize that a mechanical stretch-ECM production feedforward mechanism exists, whereby Ca²⁺ responses in FBs are triggered by mechanical stimuli, such as vessel dilation or tissue tension. This mechanism likely ensures the production of sufficient structural proteins to provide mechanical support and protection to the leptomeninges under physiological conditions.

### Fibroblast-macrophage crosstalk in chronic inflammation

Following a one-week injection of LPS, the leptomeninges exhibit chronic inflammation characterized by dense macrophage infiltration and close proximity between FBs and macrophages. FBs and macrophages are key players in the inflammatory response and subsequent tissue remodeling [40, 41]. Following multiple sclerosis and systemic inflammation, a large number of immune cells, including macrophages, directly enter the subarachnoid space from the dura mater via arachnoid cuff exit points [42], likely promoting activation and interaction between FBs and macrophages. Both activated FBs and macrophages can produce signaling molecules like retinoic acid, potentially influencing tissue recovery, while FBs also contribute to scar formation [43]. In our chronic systematic inflammation, only PiaFs display significantly weaker and less frequent spontaneous Ca²⁺ events, while that of ArachFs and PVFs remains unaffected. This subgroup-specific alteration suggests that PiaFs are strategically positioned to be most impacted by chronic inflammation. One possible explanation is the substantial influx of macrophages into the subarachnoid space—a region where they are typically sparse under physiological conditions—potentially altering PiaF function due to their proximity.

Despite the well-documented enhancement of FB-macrophage crosstalk during inflammation [36, 44, 45], and the role of LPS as a systemic inflammatory trigger, our results showed that none of the FB subgroups exhibit elevated calcium activity. Strikingly, the leptomeningeal FBs display an opposite trend, with less frequent and lower-amplitude spontaneous Ca²⁺ transients observed in PiaFs. This observation leads us to hypothesize that chronic inflammation does not inherently enhance FB calcium signaling. Instead, prolonged exposure to inflammatory mediators may downregulate calcium signaling, potentially as a protective mechanism to prevent sustained activation. Unregulated Ca²⁺ pathways could otherwise exacerbate fibrosis, as excessive calcium signaling has been linked to severe fibrotic outcomes [46]. Furthermore, chronic inflammation appears to impair locomotion-induced Ca²⁺ responses in PVFs, indicating a broader disruption of their physiological function in this state. Further investigations are needed to elucidate the mechanism underlying the altered FB Ca^2+^ signaling in this pathological condition.

### Role of TRPV4 channels in brain leptomeninges in health and disease

TRPV4 channels in the brain leptomeninges are expressed in vascular endothelial cells, FBs, and macrophages, with smaller amounts also found in vascular mural cells and astrocytes [2, 27, 47]. They are critical mediators of mechanosensation, osmosensation, and inflammation in the brain [48]. Previous studies showed that TRPV4 channels expressed in perivascular endfeet of astrocytes [49], vascular smooth muscle cells [50], and endothelial cells [51] are important regulators of neurovascular coupling. Furthermore, in pathological states, TRPV4 channels in FBs play a pivotal role in fibrosis, activated by both mechanical stimuli like ECM stiffness and stretch, and biochemical stimuli [20, 52]. TRPV4 channels in macrophages drive myofibroblast differentiation through the TGF-β pathway, mediating phagocytosis and cytokine secretion [52, 53].

In this study, we used intracortical cannula infusion of the TRPV4 channel antagonist GSK219 in awake mice. GSK219 non-selectively blocked all TRPV4 channels in the local cortical region. Following GSK219 infusion, locomotion speed was significantly faster compared to the control condition, yet the amplitude of vessel dilation remained unchanged (**Fig. 4F** versus **Fig. 5F**). This indicates that the efficiency of locomotion-induced neurovascular coupling was attenuated. The effect likely reflects TRPV4 channel blockade in vascular mural cells, astrocytes, and endothelial cells. Despite similar levels of locomotion-induced vasodilation with or without GSK219, the Ca²⁺ activity across all three FB subgroups was reduced. These findings suggest that TRPV4 channels mediate the locomotion-induced Ca²⁺ rise in leptomeningeal FBs. Interestingly, aside from a moderate reduction in Ca²⁺ amplitude in PiaFs, GSK219 did not significantly affect spontaneous FB Ca²⁺ activity. This suggests that TRPV4 channels are unlikely to mediate spontaneous Ca²⁺ signaling in PVFs and ArachFs, and that TRPV4 channels at PiaFs appear to function not as mediators, but as potentiators of spontaneous Ca^2+^ transients.

## Materials and Methods

### Animals

All procedures were approved by the Danish National Ethics Committee according to the guidelines set forth in the European Council’s Convention for the Protection of Vertebrate Animals used for Experimental and Other Scientific Purposes and are in compliance with the ARRIVE guidelines. Adult mice (3-6 months old) were housed on a 12-hour light: dark cycle with environmental enrichment. *Cdh5*-GCaMP8 transgenic mice (N = 13, 9 females, 4 males) were used for TPM experiments and three of them (2 females, 1 male) were also used for immunohistochemical experiments. A *Cdh5*-GCaMP8 negative littermate mouse (N = 1 female) was used for the TPM macrophage experiment.

### Surgery: Cranial window implantation

Mice were handled 5-10 min per day for 3 days before the surgery day to reduce the anxiety potentially led by post-surgery care. Carprofen (5mg/kg), ethiqa (3.25mg/kg) and dexamethasone (5mg/kg) were give s.c. 30-60 min before surgery. Mice were anesthetized with 1.5-2% isoflurane in 30% oxygen in air. Eyes were kept moist with ointment. Hair on the scalp was removed by a hair removal method and then rinsed with 70% ethanol. 100 μl of lidocaine was administered as local anesthesia 10 min prior to pain introduction. The scalp was incised, and the skull surface was roughed. Afterwards, a craniotomy was drilled over somatosensory cortex (3.5 mm lateral, 0.2 mm posterior to bregma; Φ= 4 mm). The exposed brain was gently washed by ice-chilled aCSF and subsequently sealed by a coverslip (Φ= 5 mm). The rim of coverslip was secured by adhesive resin cement (56857, 3M). Acrylic resin powder (64707948, Kulzer) was mixed by the solvent (66070741, Kulzer) and applied to the skull. Right after, a head plate (Model 5, Neurotar) was carefully positioned onto the dental cement mixture before solidification. After surgery, mice were injected s.c. with another 0.5 mL of saline and returned to a cage warmed by a heating pad. In the following 4 days, carprofen (same dosage as above) was given once a day as pain treatment.

### Surgery: Cannula and window implantation

The coverslip was processed with a mounted point (CA1063, MINITOR CO.) to create a space for cannula implantation. Other procedures prior to skull exposure were identical to those described above. Once the skull surface was roughened, a head plate was cemented onto it. Following the craniotomy, a small hole (approximately 0.5 mm in diameter) was drilled at the edge of the cranial window, avoiding areas with large underlying vessels. A guide cannula (C315GRL, Protech International) was then implanted into the cortex at an angle of 15-20 degrees. Subsequently, the preprocessed coverslip was sealed together with the cannula. Once secured, a dummy cannula (315DC/SPC, Protech International) was screwed onto the guide cannula.

### Surgery: Intracortical infusion

Mice were anesthetized with 1.5% isoflurane and warmed by a heating pad at 37 °C during infusion. The dummy cannula was slowly removed and replaced by an internal cannula (C315I/SPC, Protech International). TRPV4 antagonist GSK2193874 (1 μM) was infused by a combination of a syringe pump (sp100i, World Precision Instruments) and a 10 μl Hamilton syringe (7653-01, Hamilton Company) at a rate of 25 nL/min for 500 nL. Mice were imaged 20 min after anesthesia.

### Habituation

Habituation began at least one week post-surgery. For the first three days, animals were handled for 10-15 minutes daily. On day four, they were placed in the mobile homecage without head fixation for approximately 15 minutes to acclimate to the imaging environment. In the subsequent days, animals underwent head fixation in the mobile homecage, starting at 5 minutes and increasing to 60 minutes by the end, with daily increments of 10-15 minutes. Air puffs were introduced starting from the second head-fixed session.

### Vibrissa stimulation

Whisker air puffs (1 s duration, 3 Hz frequency, 120 ms pulse width) were delivered using a Pneumatic PicoPump (PV830, World Precision Instruments). The air puff nozzle was positioned above the contralateral side of the vibrissae to ensure the puffs did not contact the body. Air pressure was adjusted to the minimum level required to elicit a deflection. Whisker air puff was delivered after a 40-sec baseline recording.

### Systematic inflammation

Systematic inflammation was induced by intraperitoneal injection of lipopolysaccharide (LPS, 5mg/kg, 500X lipopolysaccharide from E. coli O111:B4; 00–4976-93, Thermofisher). Mice were given attractive food (Seeds or oats) to boost their appetite and two-photon imaging experiments were performed before and one week after LPS injection.

### Two-photon microscopy

Texas Red (2% wt/vol, molecular weight 70000, 20 μL; ThermoFisher) was retro-orbitally injected into blood plasma when mice were briefly anesthetized by isoflurane. Before waking up, mice were head-fixed onto a locomotion platform (NTR000251-06, Neurotar). Experiments were performed by a commercial two-photon microscope (FVMPE-RS, Olympus) and a 25 × 1.05 N.A. water-immersion objective (XLPLN15XWMP2, Olympus). The excitation laser was set to 920 nm. The emission signals from GCaMP8 (Cadherin 5-positive cells) and Texas Red (plasma) were filtered by band-path filters 489 - 531 nm and 601 – 657 nm, respectively. The SHG (collagen dominant) signal was excited by a 920-nm laser as well and filtered by a band-pass filter at 420 – 465 nm. A 2-dimensional x-y scan was performed to acquire a cross-section scan over the leptomeninges. The frame size along the x-axis was 1028 × 1028, and the step along the z-axis was set between 0.5 μm and 2 μm according to the size of arterioles. The temporal resolution ranged from 0.6 s to 0.9 s. The imaging was synchronized with the locomotion tracking by a built-in sensor in the Neurotar Mobile Homecage system and its software (Locomotion Tracker Version 3.0.0.65, Neurotar). During the whole process, the general behavior of animals was recorded by a webcam (Lifecam Studio, Microsoft).

### Two-photon microscopy image analysis

The imaging analytical tool was custom-made using MATLAB R2018a. To measure vessel diameters, rectangular ROIs with a width of 4 ∼ 6 μm were drawn across the center of vessel cross-section. The rectangular ROI was averaged by projection into one line for each frame, representing the profile of the vessel segment at this frame. The profile line was plotted as a two-dimensional image with the x-axis representing the number of frames. Chan-Vese segmentation or pixel intensity-based segmentation was used to delineate the vessel edge and estimate the diameter change at this ROI. To measure Ca^2+^ signals, hand-drawn ROIs with flexible contours were placed surrounding the region of FB subgroups.

In particular, we developed a specific algorithm to precisely measure PVF Ca^2+^, accounting for the altered position of PVFs due to vessel diameter change. The detailed steps are as follows: Firstly, we estimated the vessel contour in each frame, and the endothelial layer was identified by setting its upper thickness limit to 5 μm and by its characteristic morphology, which tightly enwraps the red vessel lumen. Next, vascular smooth muscle cells were identified as lying outside the endothelial layer and appearing as a black ring in our TPM. Lastly, the PVF layer was identified as a partial ring-shaped structure outside endothelial layer, and co-localized with GCaMP8 signal. For each analysis, we closely reviewed the TPM video, and tracked the identified PVF layer during the vessel diameter change. A more refined definition of the thickness at each layer may be necessary to generate a time-series dynamic ROI for PVFs accurately.

Pixels of foreground (Ca^2+^) at each frame were selected as above mean + 2×std of all pixels in the ROI. Ca^2+^ signals at each frame were calculated as average of all foreground pixel intensities at this frame. Thus, time course of Ca^2+^ waveform was plotted. After estimation a rough baseline using asymmetric least squares smoothing, we calculated mean and std of the baseline, and further normalized the Ca^2+^ waveform. Onset of both diameter and Ca^2+^ were defined as the first point above mean + 2×std of baseline. Duration of both diameter and Ca^2+^ were defined as the distance between first and last point above mean + std of baseline without falling beneath this threshold. Ca^2+^ transient frequency was defined as total number of Ca^2+^ transients per 1 minute. Percentage of Ca^2+^ active period was defined as the summation of all Ca^2+^ transient periods divided by the total recording time in this trial.

### Imaging of macrophages

LPS was administered in a wild-type mouse 7 days prior to imaging. A monoclonal F4/80 antibody conjugated with Alexa Fluor™ 488 (8 μg in 20 μl PBS, 53-4801-80, Thermo Fisher) was injected retro-orbitally 12 hours before surgery. The mouse was deeply anesthetized with xylazine (10 mg/kg) and ketamine (60 mg/kg), followed by tracheostomy and craniotomy. Texas Red was injected into the bloodstream, and the mouse was then transferred for imaging.

### Electron microscopy (EM) data access and 3D reconstruction

We used ultrastructural data from the MICrONS electron microscopy dataset [54], available at www.microns-explorer.org. The dataset includes 1 µm^3^ of a male mouse visual cortex, imaged at 4x4x40 nm. While the cortical cells and vasculature are segmented in 3D, the pial surface and leptomeninges are not. We extracted the pial surface and leptomeninges around a large pial arteriole (location in Neuroglancer: 87644, 91688, 8900) using Cloudvolume (Python) coordinates [11111:11310, 42398:46848, 39879:46901]. Segmentation was done with TRAKEM2 (FIJI), focusing on cell surfaces, nuclei, and centrioles/primary cilia. The 3D structures were visualized using Blender.

### Immunohistochemistry

Mice were deeply anesthetized with xylazine (10 mg/kg) and ketamine (60 mg/kg), then transcardially perfused with phosphate-buffered saline (PBS) followed by fixation with 4% paraformaldehyde. Brains were extracted and immersed in 4% paraformaldehyde at 4°C for 2 hours before being transferred to 30% sucrose in PBS for 48 hours. Subsequently, they were snap-frozen in isopentane and stored at −80°C until processing. Brains were sectioned at the somatosensory regions to a thickness of 50 μm using a cryostat. Sections were rinsed with PBS for 5 minutes, three times, before simultaneous permeabilization and blocking with 0.5% Triton X-100 and 1% BSA in PBS at 4°C overnight. The sections were then incubated with primary antibodies: anti-GFP (13970, Abcam), anti-mouse CD31 (553371, BD Bioscience), and anti-mouse PDGFRα (AF1062, R&D systems) for 30 hours at 4°C, followed by rinsing and incubation with secondary antibodies (1:500). Afterward, they were immersed in Hoechst (1:4000) for 7 minutes. Following three 5-minute washes with PBS, sections were mounted on glass slides using SlowFade Diamond Antifade Mountant (S36963, Invitrogen). Brain samples were imaged using a Zeiss LSM 980 confocal microscope.

### Statistical analysis

Datasets are presented as the mean ± s.t.d. with individual data points. The normality of data was assessed using Shapiro–Wilk and graphical tests. If the data were not normally distributed, they were log-transformed and retested for normality. Linear mixed-effect model analysis was used. FB subgroups (ArachF, PiaF, PVF) and treatment (control, GSK219, LPS) were defined as the fixed effect, whereas the mouse ID, imaging location, and imaging date were included as random effects. Significant differences were estimated by likelihood ratio tests of the linear mixed-effect model with the fixed effect in question against a model without the fixed effect. Tukey–Kramer’s post hoc test was used for pairwise comparisons between elements in the fixed-effect subgroup. All statistical analyses were performed using Rstudio (version 3.4.4, packages lme4 and multcomp).

## Data and source code availability

Source data are provided with this paper (10.5281/zenodo.15382814). All other data are available upon request. The customized MATLAB code is uploaded to Github: https://github.com/ChangsiCai/Matlab-code-for-Fibroblast-manuscript

## Acknowledgements

We thank Assoc. Prof. Carmelo Bellardita in the Department of Neuroscience, University of Copenhagen, who discussed with us and provided us with technology to visualize macrophages *in vivo*. This study was supported by the Lundbeck Foundation (R345-2020-1440 and R436-2023-1125), the Danish Medical Research Council (1133-00016B), the Novo Nordisk Foundation (NNF0064289) and a Nordea Foundation Grant to the Center for Healthy Aging, the Alice Brenaa Foundation, Augustinus Foundation, Carl og Ellen Hertz Familielegat, A. P. Møller Foundation, Helsefonden, Dagmar Marshall Fond, Oda og Hans Svenningsens Fond and Torben og Alice Frimodts Fond.

## Authorship contribution statement

Conceptualization: CH, CC; Data curation: CH, SG; Formal analysis: CH, SG, CC; Funding acquisition: CC; Investigation: CH, SG, LT, XZ; Methodology: CH, SG, LT, XZ, KK, AD, NZS; Resources: CH, SG, CC; Software: CH, SG, CC; Supervision: CC; Writing: CH, SG, CC.

## Supplementary figure texts

**Supplementary Figure 1.**
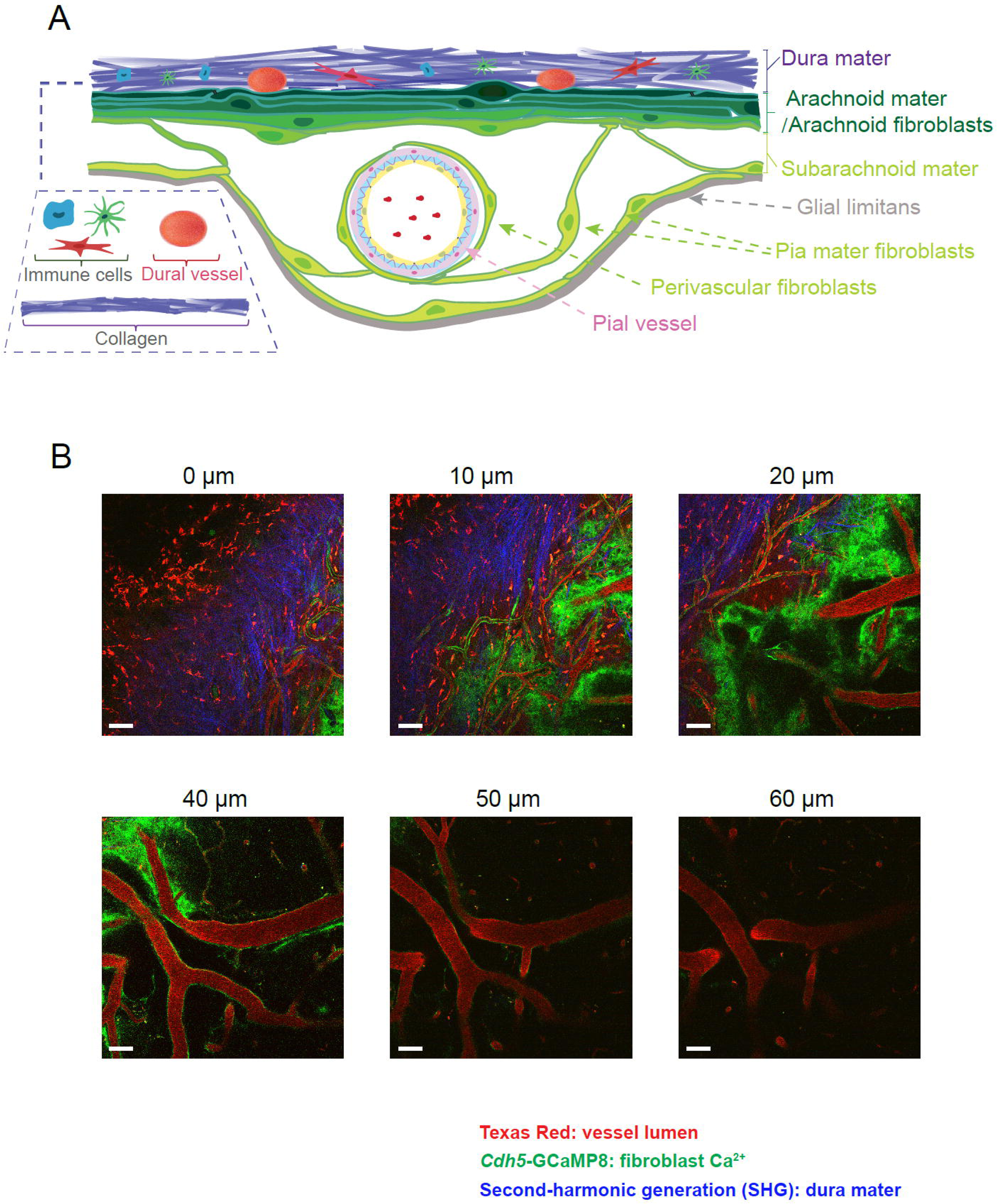
Diagram and two-photon microscopy of leptomeningeal fibroblasts. (**A**) Diagram of locations and morphologies of each leptomeningeal FB subgroups. (**B**) x-y plane TPM images at different cortical depths. Red: i.v. injection of Texas Red for vessel lumen. Green: *Cdh5*-GCaMP8. Blue: second harmonic generation (SHG) for collagen. Scale bar: 50 μm.

**Supplementary Figure 2.**
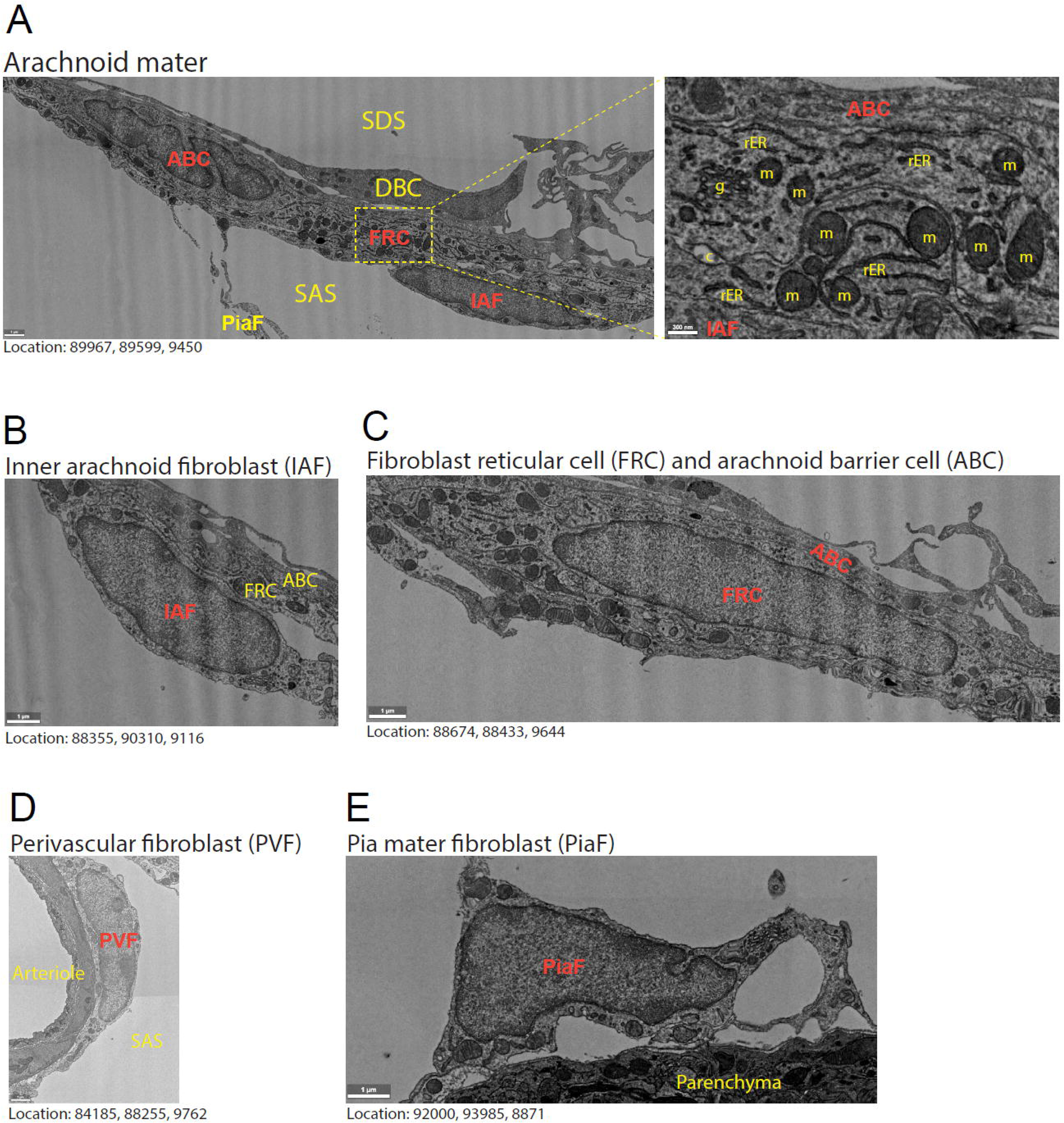
Ultrastructures of leptomeningeal FB subgroups based on EM data. (**A**) Showing the ultrastructure of cells forming the arachnoid mater separating the subarachnoid space (SAS) from the subdural space (SDS). Processes of pia mater FBs (PiaF) or inner arachnoid FBs (IAF) form trabeculae that cross the SAS. The IAFs form the inner lining of the arachnoid mater, adhering to the FB reticular cells (FRC). Morphologically, it is difficult to distinguish PiaFs, IAFs and perivascular FBs (PVFs), so their identities are based on the location of their nuclei. FRCs are distinguishable by their high content of rough endoplasmic reticulum (rER) and a large mitochondrial (m) network. Collagen (c) strands form a matrix of intercellular channels in the FRC layer that is connected with the subpial collagen matrix via larger collagen fibers. Arachnoid barrier cells (ABCs) cover a large area and often have more than one nucleus (this ABC has 2 nuclei). They have thin processes that overlap other ABC processes tightly and have no collagen strands or channels crossing them, forming a diffusion barrier. The FRCs adhere to ABCs. In some places, remnants of the removed dura in the form of dural border cells (DBC) loosely attach to the ABCs. Golgi apparatus (g). Scale bar: 1 μm. Inset: 300 nm. (**B-E**) Showing close-up examples of the different cell types of the leptomeninges. Scale bar: 1 μm. Notice how PVF, PiaF, and IAF which line the SAS have a similar morphology, while FRCs have more rER and mitochondria. Locations indicate coordinates in the dataset accessible at www.microns-explorer.org

**Supplementary Figure 3.**
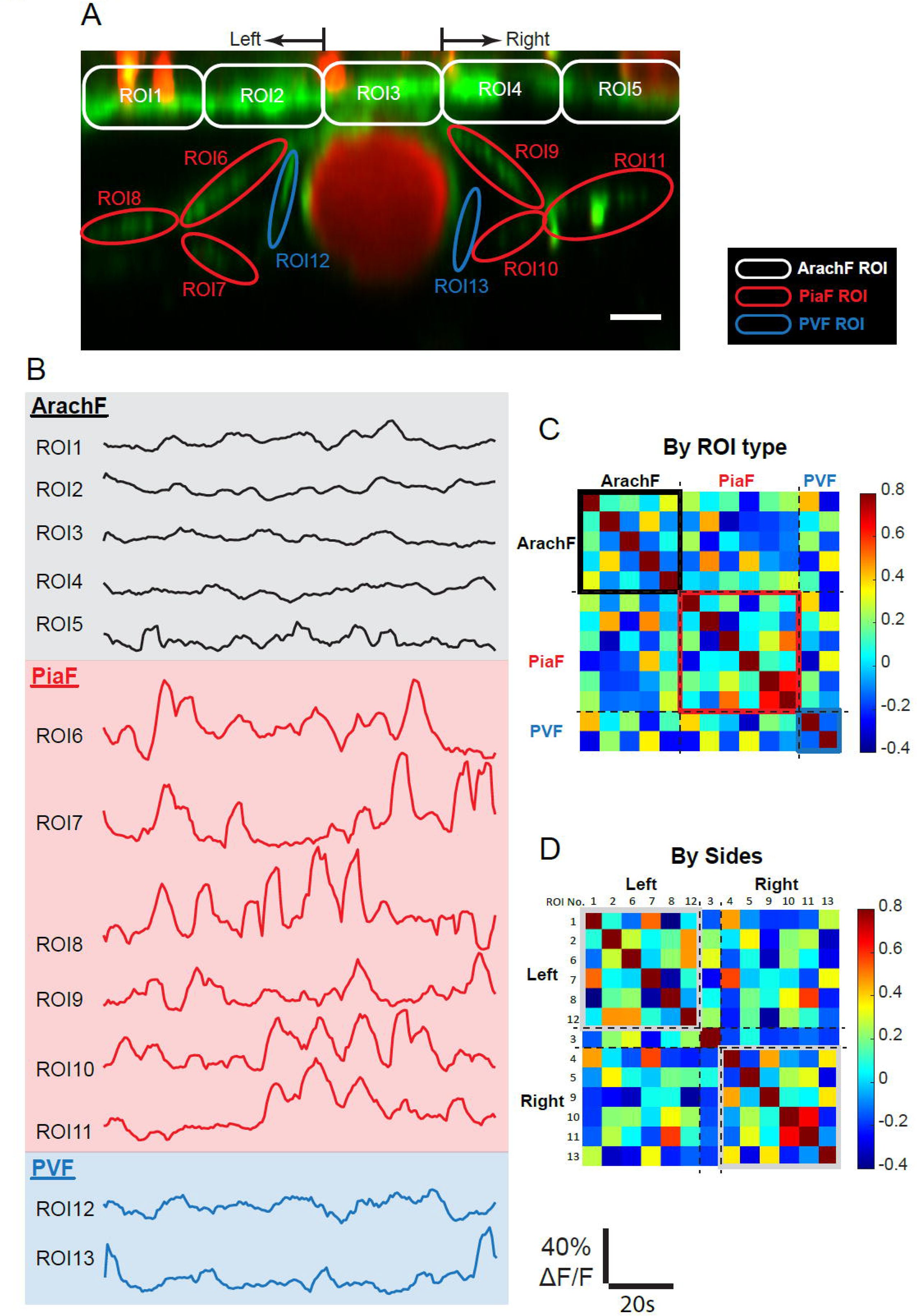
Uncorrelated fibroblast Ca^2+^ activities in a representative two-photon recording. (**A**) x-z plane TPM image. Red channel: Texas Red for vessel lumen. Green channel: Cadherin5-GCaMP8. ROIs at each fibroblast (FB) subgroup are marked with different colors and sequential numbers. Scale bar: 10 μm. (**B**) Measured Ca^2+^ activities at each ROI. (**C**) Heat map of correlation coefficient sorted by FB subgroups. (**D**) Heat map of correlation coefficient sorted by ROI spatial proximity.

**Supplementary Figure 4.**
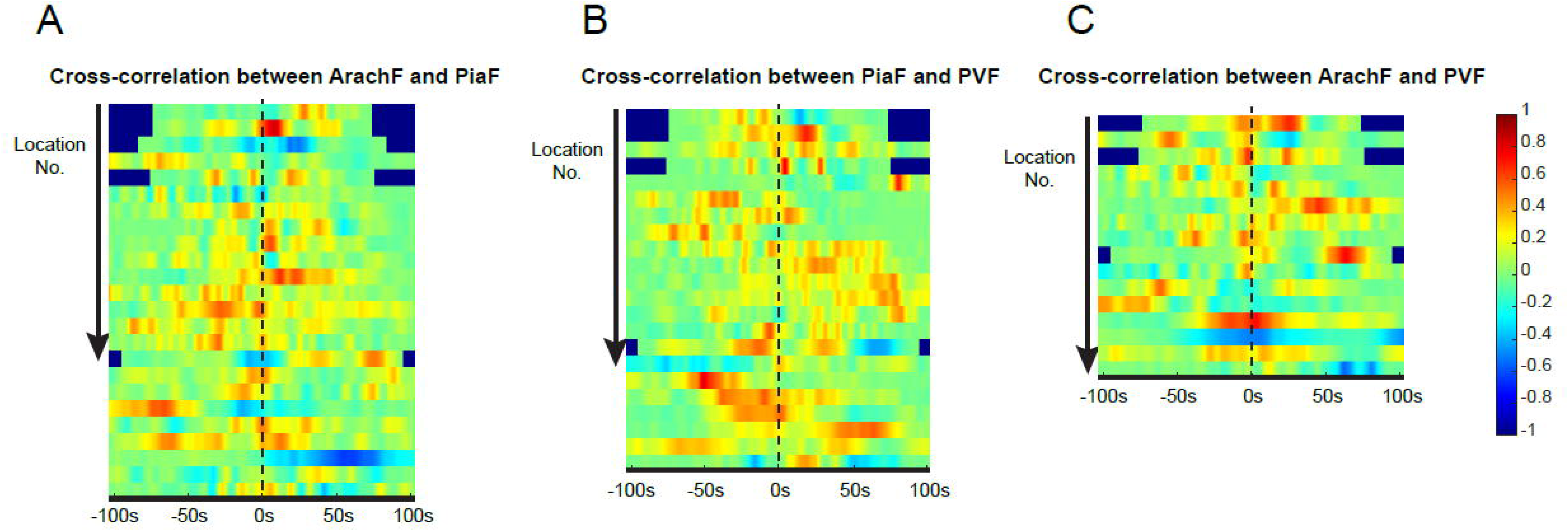
Uncorrelated fibroblast Ca^2+^ activities by paired fibroblast subgroups. (**A**) Heat map of cross-correlation between ArachF and PiaF. The X-axis represents the time delay of the cross-correlation of each pair of traces. The y-axis represents location number. (**B**) Heat map of cross-correlation between PiaF and PVF. (**C**) Heat map of cross-correlation between ArachF and PVF.

**Supplementary Figure 5.**
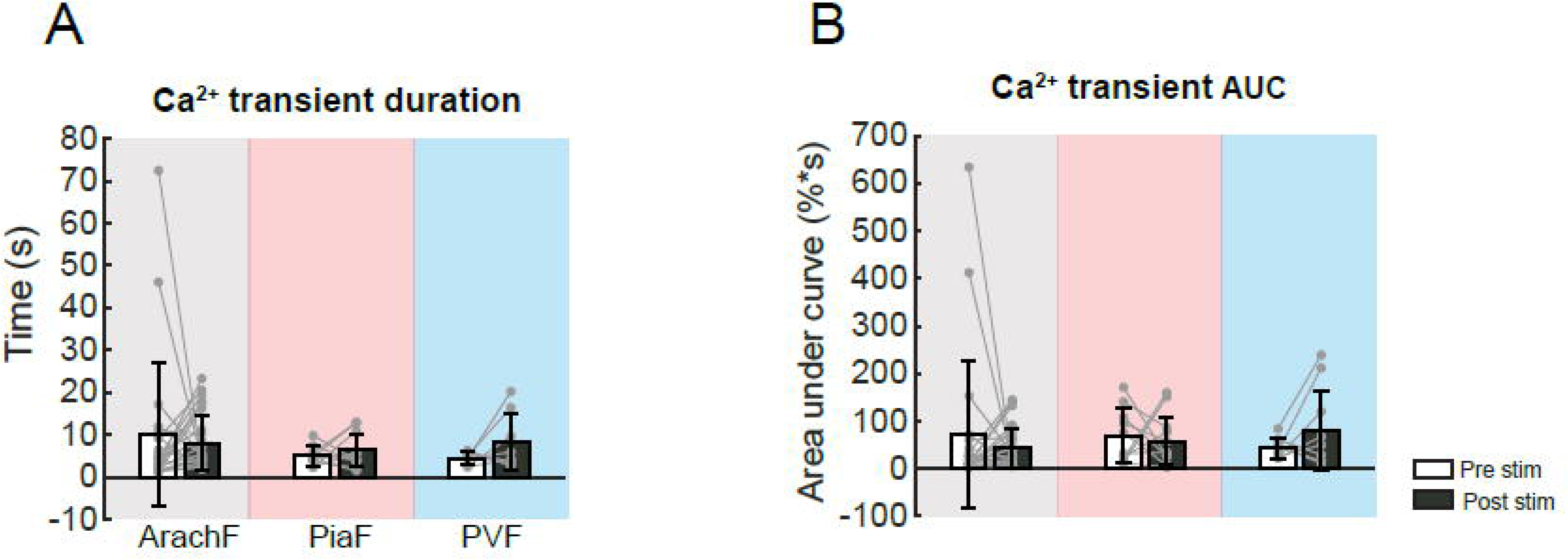
Comparison of fibroblast Ca^2+^ transients before and after whisker air puff. (**A**) Comparison of Ca^2+^ transient duration before and after air-puff stimulation. (**B**) Comparison of Ca^2+^ transient area under the curve (AUC) before and after air-puff stimulation. (ArachF: N = 4 mice, n = 9 locations, n′ = 29 trials; PiaF: N = 3 mice, n = 8 locations, n′ = 13 trials; PVF: N = 4 mice, n = 7 locations, n′ = 8 trials.) Bar plots are presented as the mean ± s.t.d. Linear mixed-effect models were used, followed by Tukey post hoc tests for pairwise comparisons. * p < 0.05, ** p < 0.01, *** p < 0.001.

**Supplementary Figure 6.**
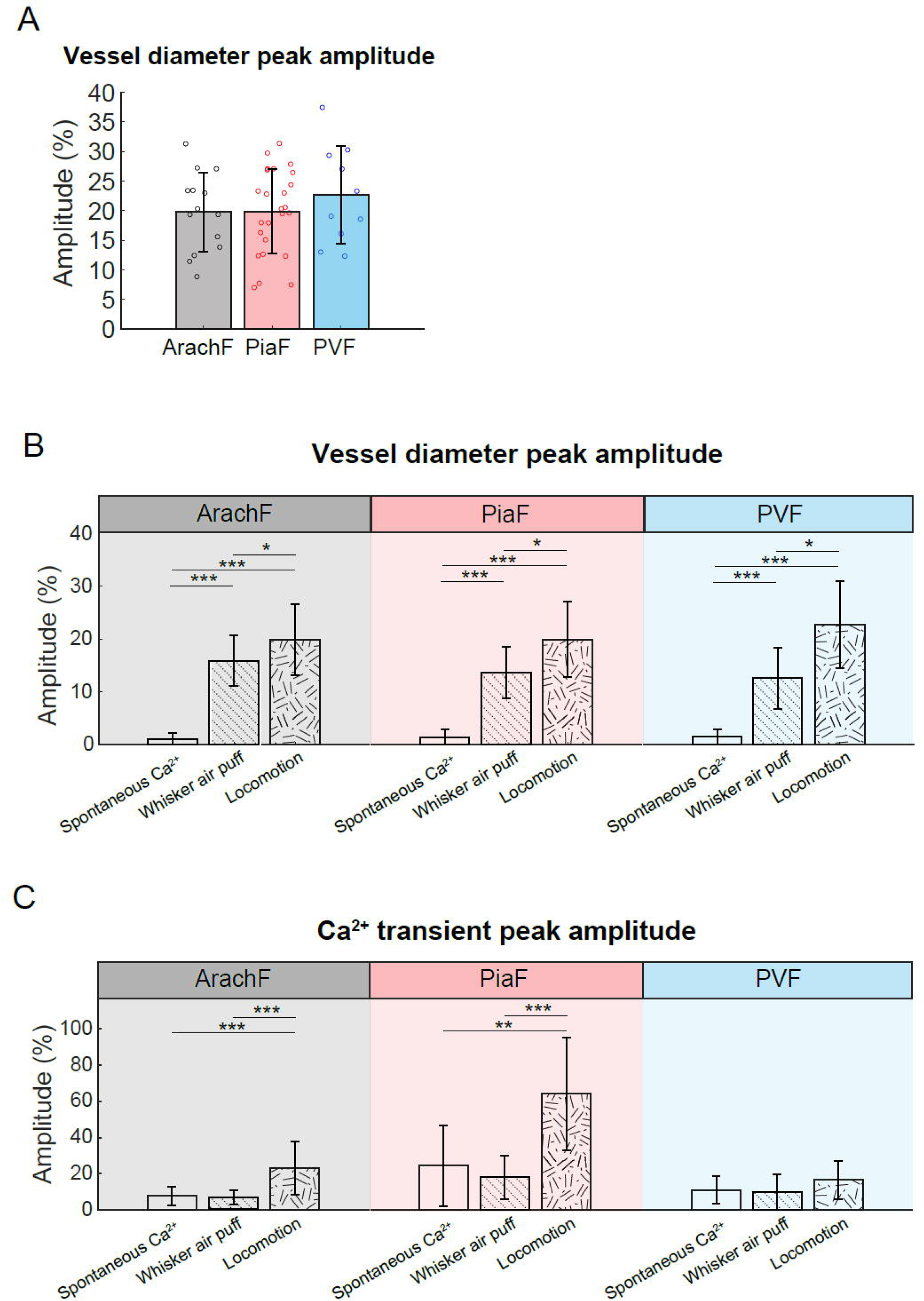
Summary of the Ca^2+^ profiles and co-localized vessel diameter change at three fibroblast subgroups. (**A**) Comparison of locomotion-induced vessel diameter change at each fibroblast (FB) subgroup, focusing on peak amplitude. (ArachF: N = 5 mice, n = 7 locations, n′ = 35 trials; PiaF: N = 6 mice, n = 9 locations, n′ = 25 trials; PVF: N = 4 mice, n = 8 locations, n′ = 10 trials.) (**B-C**) Comparison across three scenarios: spontaneous Ca^2+^, whisker air puff, and locomotion. Peak amplitude of vessel diameter (B) and Ca^2+^ transients at each FB subgroups are compared. Bar plots are presented as the mean ± s.t.d. Linear mixed-effect models were used, followed by Tukey post hoc tests for pairwise comparisons. * p < 0.05, ** p < 0.01, *** p < 0.001.

**Supplementary Figure 7.**
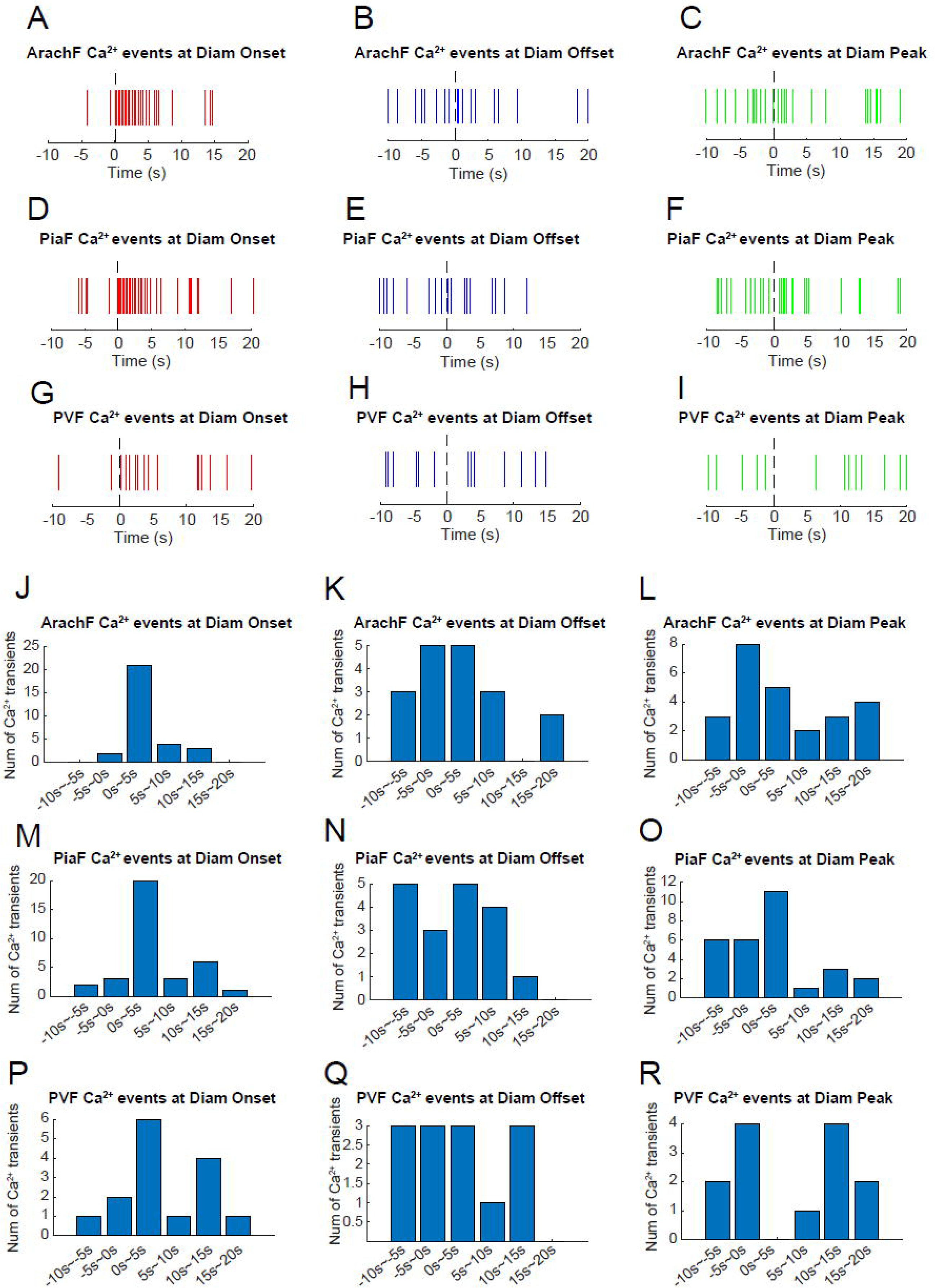
Locomotion-induced fibroblast Ca^2+^ transients occur predominantly with vessel diameter onset. (**A-I**) The onset of each Ca^2+^ transient is digitized as a vertical line, and overlaid at different time points of vessel diameter change, i.e. vasodilataion onset, peak and offset time. Each row represents each FB subgroup, with ArachF, PiaF and PVF respectively. Each column represents different time points of vasodilation, with onset, peak time and offset respectively. (**J-R**) are summarized bar plots based on (A-I). The height of bars represent the number of Ca^2+^ transients, x-axis represents the multiple 5-second-width windows before and after each vasodilation time point. 0s are the moment of occurrence of onset, peak, and offset of vasodilation, respectively. (ArachF: N = 5 mice, n = 7 locations, n′ = 35 trials; PiaF: N = 6 mice, n = 9 locations, n′ = 25 trials; PVF: N = 4 mice, n = 8 locations, n′ = 10 trials.)

**Supplementary Figure 8.**
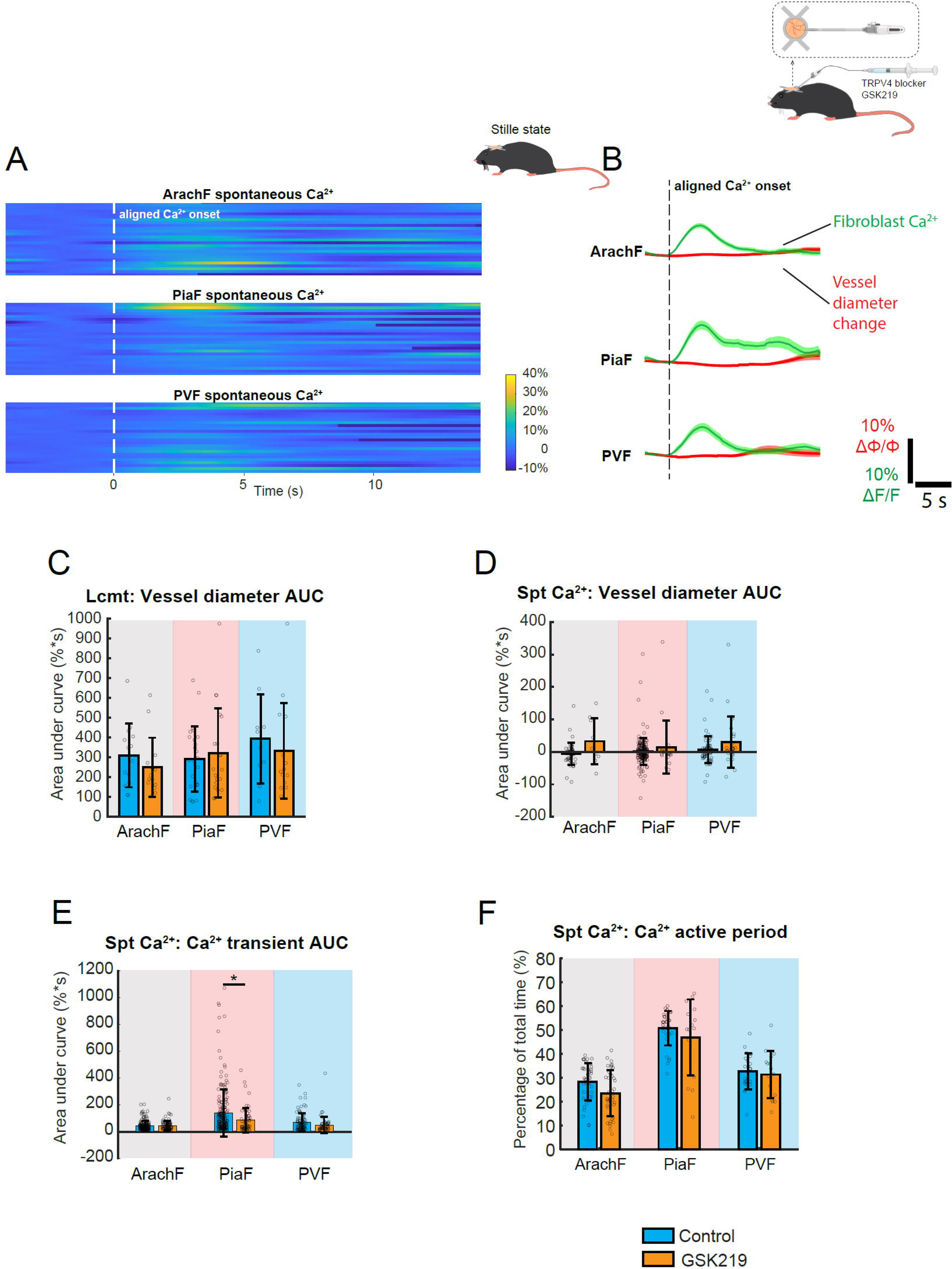
TRPV4 channel blocker GSK219 alters fibroblast Ca^2+^ profiles. (**A**) After administration of GSK219, heat maps of spontaneous Ca^2+^ transients at each FB subgroup, by aligning Ca^2+^ onsets. x-axis: time. Y-axis: trace number. (**B**) Overlaid Ca^2+^ and vessel diameter traces for each FB subgroups after GSK219. The shadow represents s.e.m. of all traces. (**C**) Comparison of locomotion-induced Ca^2+^ responses without or with intracortical infusion of GSK219, focusing on the area under the curve of vessel diameter change. (**D-F**) Comparison of spontaneous Ca^2+^ transients without or with intracortical infusion of GSK219, focusing on (D) area under the curve of vessel diameter change, (E) area under the curve of Ca^2+^ transients, and (F) Ca^2+^ transient active period. (**GSK219 spontaneous Ca^2+^** -ArachF: N = 5 mice, n = 11 locations, n′ = 93 events; PiaF: N = 5 mice, n = 11 locations, n′ = 63 events; PVF: N = 5 mice, n = 11 locations, n′ = 61 events.) (**GSK219 locomotion -** ArachF: N = 4 mice, n = 11 locations, n′ = 38 trials; PiaF: N = 3 mice, n = 9 locations, n′ = 21 trials; PVF: N = 3 mice, n = 10 locations, n′ = 14 trials.) Linear mixed-effect models were used, followed by Tukey post hoc tests for pairwise comparisons. * p < 0.05, ** p < 0.01, *** p < 0.001.

**Supplementary Figure 9.**
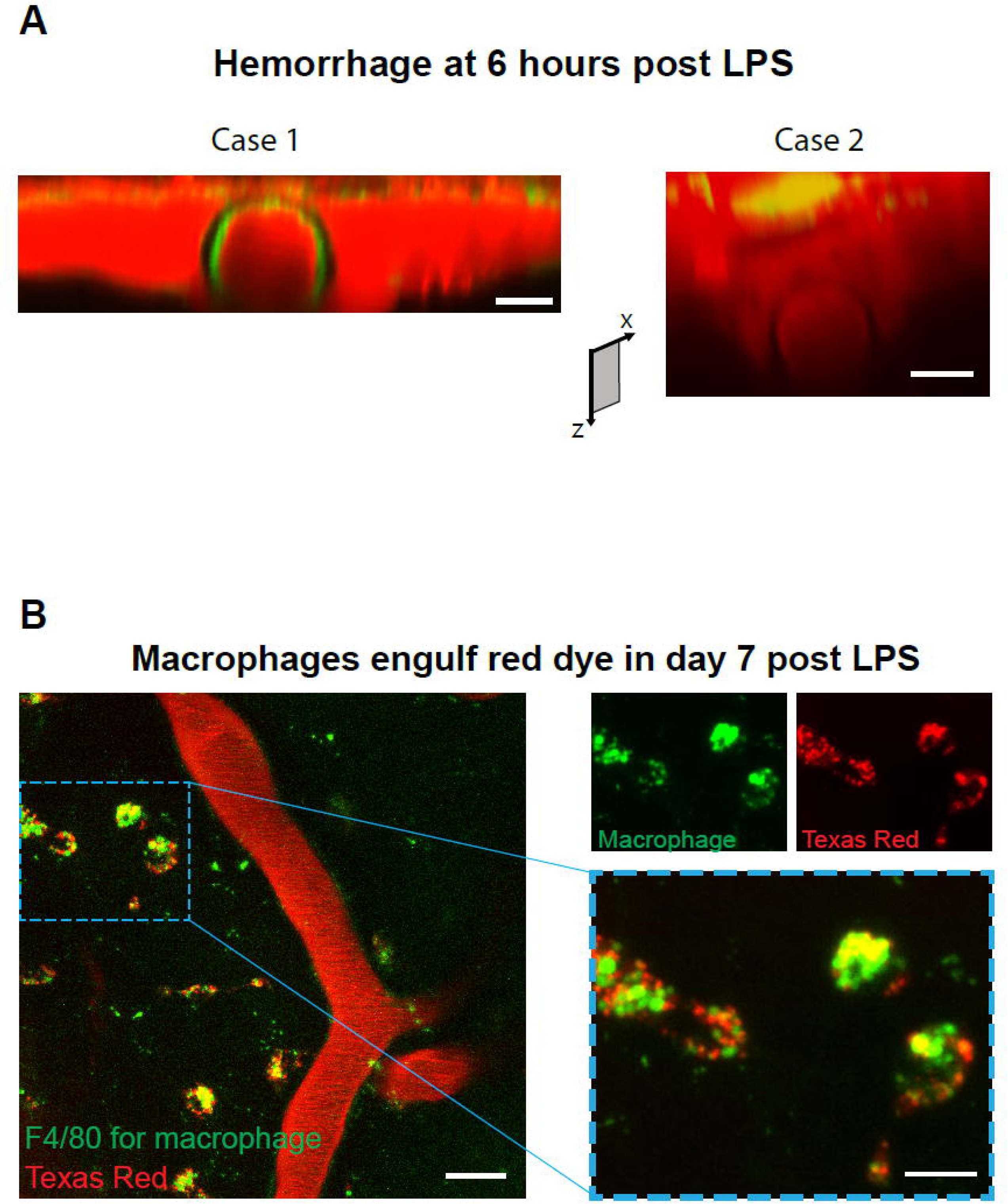
LPS-associated systemic inflammation leads to disruption of the blood-brain barrier in the leptomeninges. (**A**) Six hours after i.p. injection of LPS, almost all the examined mice experienced dye leakage (red cloud) in the leptomeninges, likely due to the disrupted blood-brain barrier by systemic inflammation. This situation severely affects the imaging quality of FB Ca^2+^ signals. x-z plane two-photon image. Scale bar: 20 μm. (**B**) The leaked red dye is taken up by leptomeningeal macrophages. Green: F4/80-Alexa488 antibody to stain macrophages. Texas red: red vessel lumen dye. Left panel: large-scale x-y TPM image. Scale bar: 25 μm. Right panel: zoomed-in image of a local region. Red and green channels are co-localized but not overlaid. Scale bar: 10 μm.

**Supplementary Figure 10.**
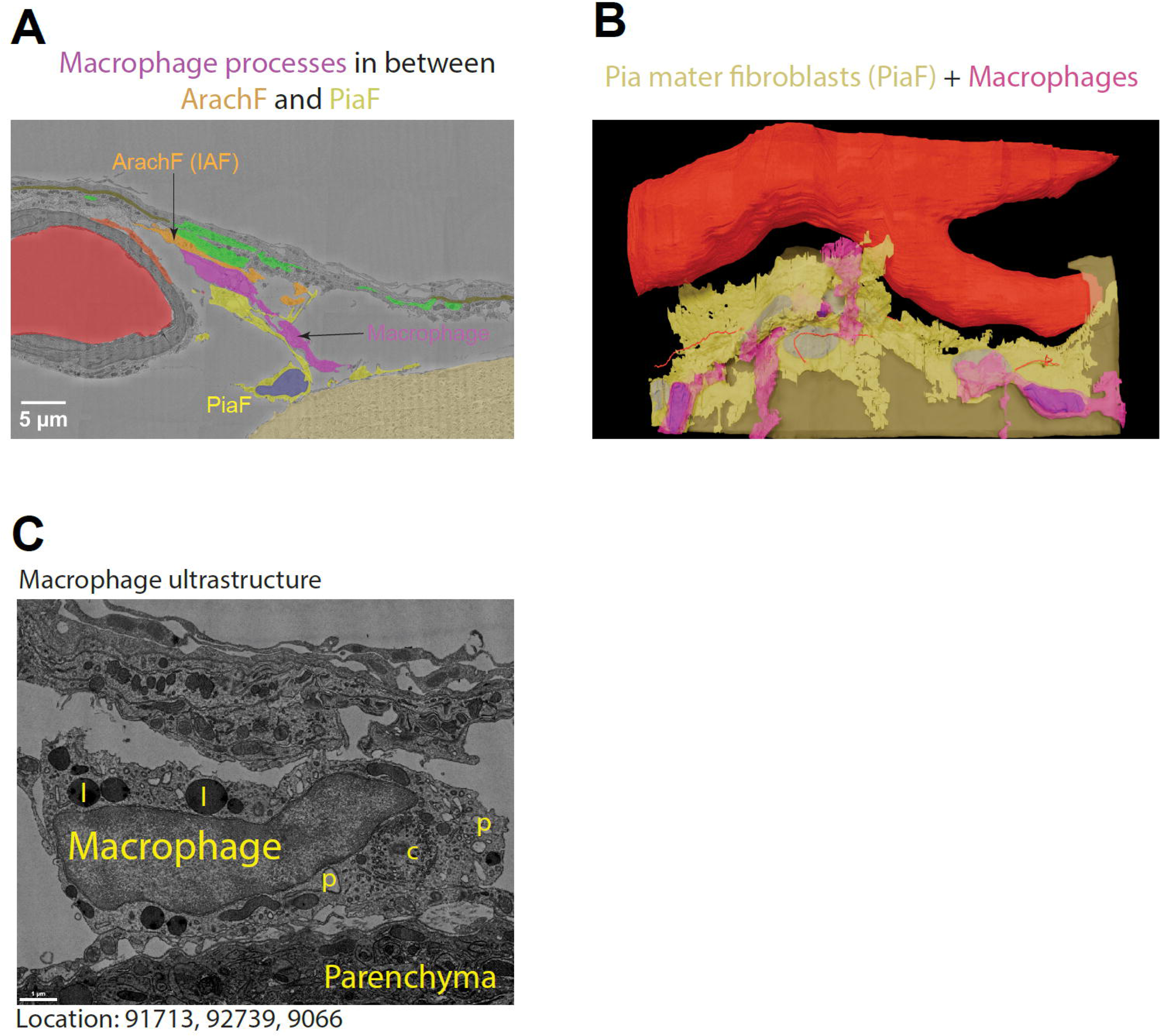
Ultrastructure of leptomeningeal fibroblasts and macrophages. (**A**) The ultrastructure data shows how macrophage processes can be localized between the ArachF and PiaF processes. (**B**) 3D segmentation shows 3 macrophages that are either on top of the PiaFs or in between them. (**C**) Macrophages are recognized by their many lysosomes (l) and phagosomes (p), but also by the absence of primary cilia and the centrioles (c) located in the center of the Golgi apparatus. Location indicates coordinates in the dataset accessible at www.microns-explorer.org.

**Supplementary Figure 11.**
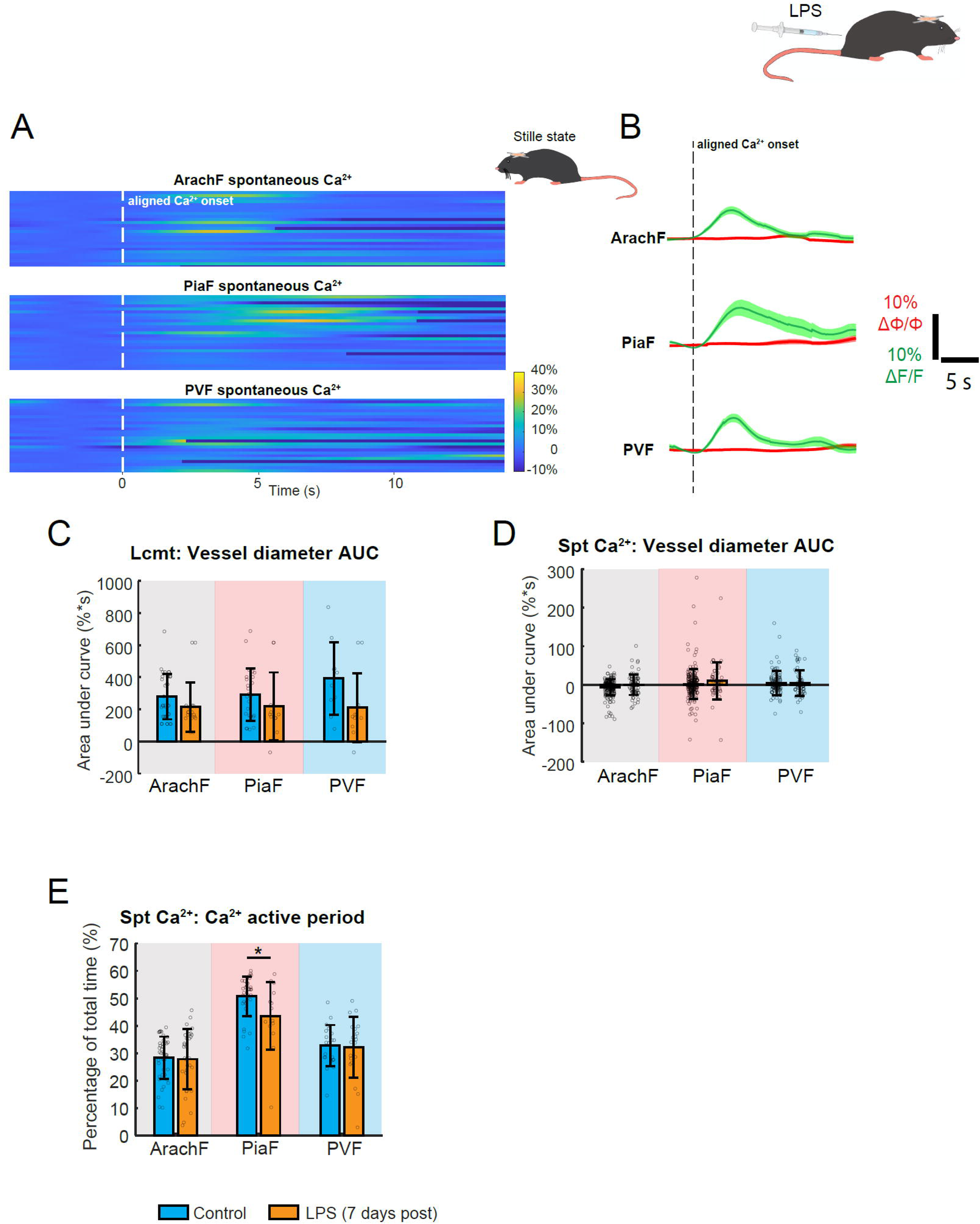
LPS-associated systematic inflammation alters fibroblast Ca^2+^ profiles. (**A**) Seven days after administration of LPS, heat maps of spontaneous Ca^2+^ transients at each FB subgroup were aligned by Ca^2+^ onsets. x-axis: time. Y-axis: trace number. (**B**) Overlaid Ca^2+^ and vessel diameter traces for each FB subgroups after LPS. The shadow represents s.e.m. of all traces. (**C**) Comparison of locomotion-induced Ca^2+^ responses without or with LPS, focusing on the area under the curve of vessel diameter change. (**D-E**) Comparison of spontaneous Ca^2+^ transients without or with LPS, focusing on (D) area under the curve of vessel diameter change and (E) Ca^2+^ transient active period. (**LPS spontaneous Ca^2+^** -ArachF: N = 3 mice, n = 9 locations, n′ = 80 events; PiaF: N = 3 mice, n = 8 locations, n′ = 46 events; PVF: N = 3 mice, n = 11 locations, n′ = 62 events.) (**LPS locomotion -** ArachF: N = 3 mice, n = 7 locations, n′ = 18 trials; PiaF: N = 3 mice, n = 7 locations, n′ = 11 trials; PVF: N = 3 mice, n = 8 locations, n′ = 11 trials.) Linear mixed-effect models were used, followed by Tukey post hoc tests for pairwise comparisons. * p < 0.05, ** p < 0.01, *** p < 0.001.

## Video legends

**Supplementary Video 1.** Cascade of 2D ultrastructures of ArachF, PiaF and PVF using the MICrONS dataset of high-resolution electron microscopy.

**Supplementary Video 2.** 3D reconstruction of a local leptomeningeal fibroblast ultrastructure from MICrONS dataset. Each subgroup of fibroblasts is added sequentially.

**Supplementary Video 3.** Time-series movie of leptomeningeal fibroblast Ca^2+^ at x-z plane. The awake mouse is in a stilled state.

**Supplementary Video 4.** Time-series movie of leptomeningeal fibroblast Ca^2+^ at x-z plane. The awake mouse has spontaneous locomotion.

**Supplementary Video 5.** 3D video of macrophage infiltration.

